# Potentiating CAR-T bystander killing by enhanced Fas/FasL signaling mitigates antigen escape in heterogeneous tumors

**DOI:** 10.1101/2025.09.22.677496

**Authors:** Matthew J. Lin, Joanna K. Chorazeczewski, Gvantsa Pantsulaia, Alan Cooper, Moah Sohn, Sidorela Reci, Jaime Mateus-Tique, Nicole H. Hirsh, Xinping Xie, Ivan Odak, Rudra Dutta, Daniel Charytonowicz, Ranjan Upadhyay, Brian D. Brown, Miriam Merad, Morgan Huse, Justin Kline, Joshua D. Brody

## Abstract

Antigen (Ag) escape is a frequent mechanism of relapse after CAR-T therapy, even though only ∼1% of leukemic and ∼0.1% of lymphoma cells are Ag⁻ at baseline. In this study, we modeled extreme Ag heterogeneity (>20%) to define how Fas/FasL-dependent bystander killing contributes to tumor clearance. Across patient cohorts, Fas expression predicted survival after CD19 CAR-T therapy, particularly in CD19-low disease. In both murine and human systems, Fas-dependent bystander killing required Ag stimulation and cell contact, operated within a defined therapeutic window, and could eradicate large fractions of Ag⁻ tumors *in vivo*. Pharmacologic potentiation with inhibitor of apoptosis protein antagonists or genetic stabilization of CAR-T membrane-bound FasL enhanced bystander killing but simultaneously induced CD4⁺ T cell fratricide, which was rescued by CAR-T *Fas* knockout. Importantly, Fas sensitization also enabled bispecific antibody-redirected T cells to mediate bystander killing in resistant tumors. Finally, targeting tumor-associated macrophages triggered Fas-dependent clearance of neighboring tumor cells. These findings establish Fas-mediated bystander killing as a generalizable and therapeutically actionable axis to prevent Ag escape and broaden the scope of targeted T cell therapies.

**Graphical Abstract:** 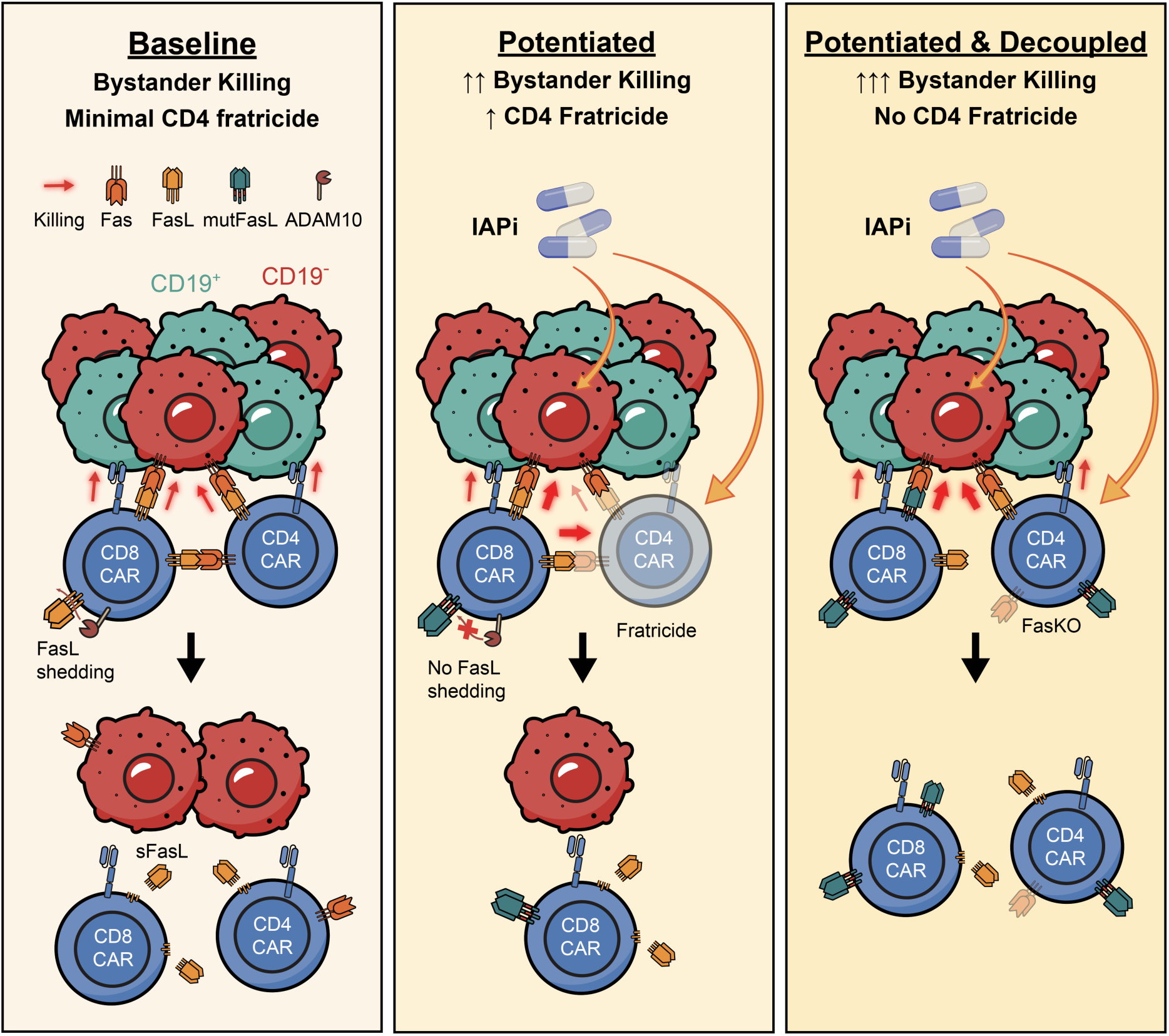

**Fas-mediated bystander killing can be potentiated by IAP inhibition and FasL stabilization, and further improved by decoupling CD4⁺ T cell fratricide via Fas knockout.**

Schematic representation of CAR-T bystander killing across three mechanistic states. **Left (Baseline):** Antigen-positive (CD19⁺) tumor cells activate CAR-T cells, leading to FasL-mediated cytotoxicity of neighboring antigen-negative (CD19⁻) tumor cells via membrane-bound FasL. Limited FasL expression and ADAM10-mediated shedding constrain bystander killing, and CD4⁺ T cells are largely spared. **Middle (Potentiated):** Enhancing Fas signaling through IAP inhibition (IAPi), increasing FasL expression, and reducing FasL shedding result in improved bystander killing but also increased CD4⁺ T cell fratricide. **Right (Potentiated & Decoupled):** Combining IAPi with FasL stabilization and CAR-T Fas knockout (FasKO) augments bystander killing while eliminating CD4⁺ T cell fratricide.

## Introduction

While chimeric antigen receptor T cell (CAR-T) therapies have dramatically improved clinical outcomes in patients with hematologic malignancies, the majority of CAR-T– treated patients do not achieve a durable remission. Among numerous causes of post-CAR-T relapse, a frequent and well-described mechanism is antigen (Ag) escape, whereby cancer cells with downregulated or absent expression of target Ag evade elimination and continue to grow.^1–4^ This phenomenon has been described in over 60% of patients with relapsed B acute lymphoblastic leukemia (B-ALL) and nearly 30% of diffuse large B cell lymphoma (DLBCL) cases after αCD19 CAR-T therapy,^5,6^ as well as patients with multiple myeloma and glioblastoma receiving CAR-T therapy.^7–10^ Ag escape also undermines responses to T-cell redirecting therapies (TRT) such as bispecific antibodies (BsAb), with reported escape rates as high as 67-88%.^11–13^ To overcome this challenge, strategies such as dual or sequential targeting of multiple Ag, logic-gated CAR-T, and prime-boost approaches have been developed.^14–19^ Though these therapies show promise, they may be limited by concurrent Ag loss (such as CD19 and CD22)^20^ or the lack of secondary targetable Ag, leaving a critical need for alternative approaches to prevent relapse.

One such mechanism is “bystander killing,” whereby Ag-specific T cells eliminate Ag^+^ cells and adjacent Ag⁻ cancer cells. This phenomenon has been linked to multiple pathways, including IFNγ,^21–27^ TNFα,^28–30^ NGK2D,^31^ and death receptor pathways such as Fas and TRAIL.^32,33^ Among these, Fas-Fas ligand (FasL) signaling plays a prominent role in direct T-cell–mediated cytotoxicity and has been shown to facilitate bystander killing.^34–36^ Our group and others have shown that Fas signaling contributes to bystander killing in both murine and human models of lymphoma.^37,38^ However, the precise dynamics, spatial constraints, and therapeutic potential of Fas-dependent bystander killing remain incompletely understood.

Despite the recognition of bystander killing as a phenomenon, its underlying dynamics, therapeutic window, and translational potential remain incompletely understood. In particular, whether Fas signaling can be leveraged to eliminate pre-existing Ag⁻ subclones and thereby prevent relapse has not been systematically explored. Here, we investigate Fas-mediated bystander killing as a determinant of CAR T efficacy in antigen-heterogeneous tumors and evaluate strategies to pharmacologically and genetically potentiate this pathway.

## Results

### FAS expression predicts better CAR-T survival outcomes in patients with low *CD19* expression

Given the contribution of a tumoral death-receptor signature (DRS) in predicting survival outcomes among B-ALL patients treated with CAR-T,^32^ we applied the same analysis to pre-treatment biopsies from a cohort of DLBCL patients receiving CAR-T and observed markedly improved survival in DRS^high^ patients (Figure 1A). Multivariate analysis of DRS component genes revealed that only *TNFRSF9* (41BB) and *FAS* contributed significantly to the prognostic ability of the DRS (Figure 1B), though their expression levels correlated with those of other downstream signaling components included in the DRS (Supplemental Figure S1A). As *TNFRSF9* transcripts in DLBCL signify T-cell infiltration,^39^ which has already been shown to predict CAR-T patient outcomes,^40^ we focused exclusively on tumor cell-intrinsic factors. Indeed, when removing *FAS* from the DRS, statistically significant survival differences between DRS^high^ and DRS^low^ patients were lost (Figure 1C).

**Figure 1.**
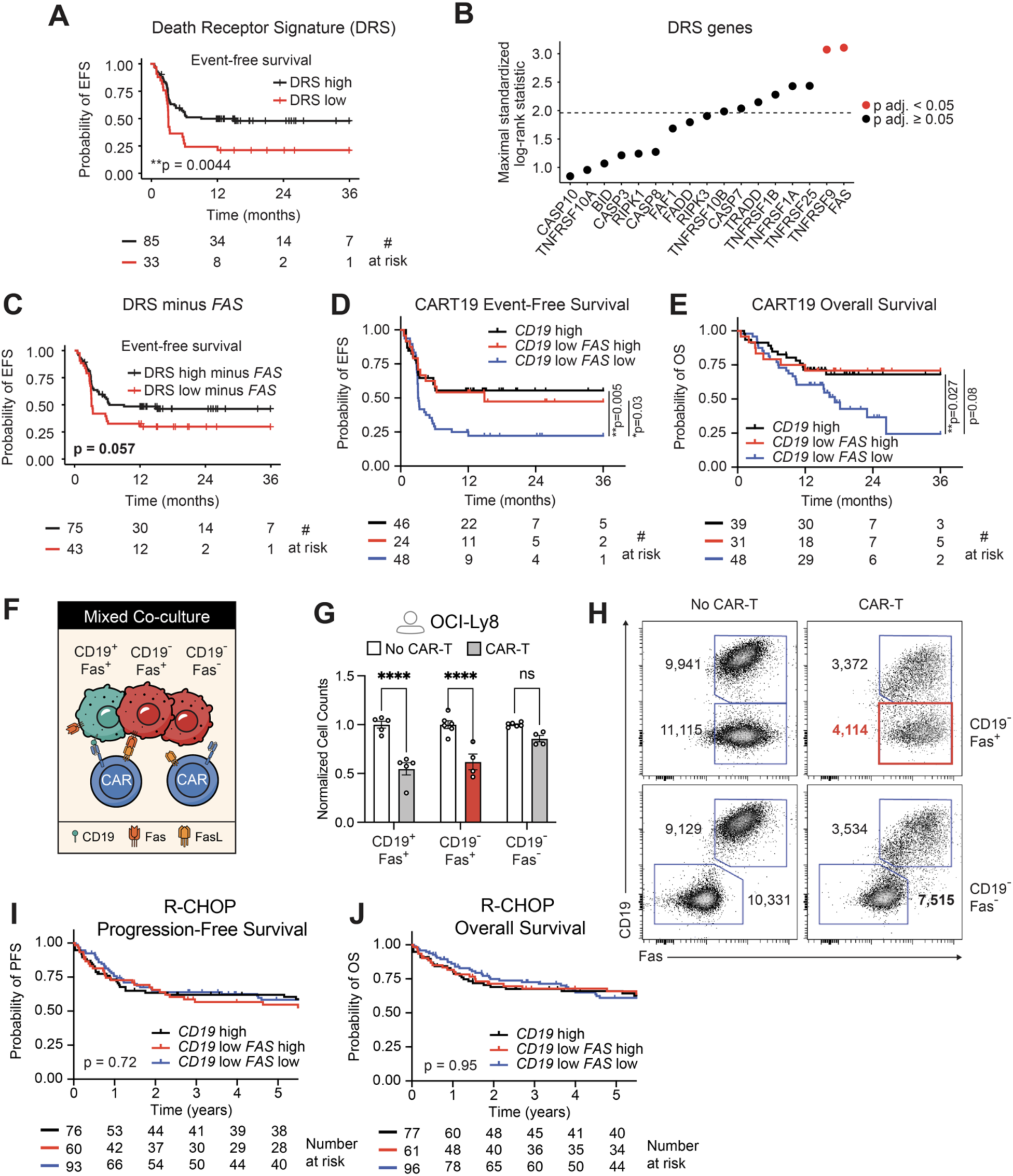
*FAS* expression correlates with survival in *CD19*-low DLBCL patients receiving CAR-T therapy and mediates Fas-dependent bystander killing in co-culture models. **(A)** Event-free survival (EFS) in DLBCL patients receiving CAR-T therapy stratified by death receptor signature (DRS). **(B)** Univariate maximal log-rank statistic for each DRS gene, reflecting the optimal cut-point for EFS, ranked by maximal test statistic. The dotted line indicates the z-score corresponding to p=0.05. Red dots denote genes significant after BH-adjustment for multiple testing. **(C)** EFS in DLBCL patients receiving CAR-T therapy stratified by DRS recalculated with *FAS* expression excluded. **(D)** EFS and **(E)** Overall survival in DLBCL patients receiving CAR-T therapy, stratified by *CD19* and *FAS* expression. Optimal cut-points (*CD19*, 60.7%; 66.7%, *FAS*) were determined using the maxstat algorithm. Log-rank tests, unadjusted. **(F)** Schematic of mixed co-culture containing equal proportions of CD19^+^, CD19⁻Fas^+^, and CD19⁻Fas⁻ tumor cells and CAR-T cells. **(G)** Normalized viable tumor cell counts 4 hours after co-culture. Dots indicate replicates, bars show means ± s.e.m. Statistical significance assessed by two-way ANOVA with Sidak correction. ns, not significant; ****p<0.0001. **(H)** Representative absolute counts of surviving CD19 and Fas-expressing tumor cell populations after co-culture. Bystander killing of CD19⁻Fas^+^ cells is highlighted by a red box. **(I)** Progression-free survival and **(J)** Overall survival in DLBCL patients receiving R-CHOP therapy from the NCI dataset. Survival differences assessed by log-rank test, unadjusted.

Because our prior transgenic T-cell studies showed that Fas signaling is essential for clearance of Ag^⁻/low^ more than Ag^+^ tumor cells, we assessed the impact of *FAS* on clinical outcomes of CAR-T patients with pre-treatment *CD19*^low^ tumors. Surprisingly, *FAS* expression was also highly predictive of improved event-free and overall survival of patients with *CD19*^low^ tumors (Figures 1D-E, Supplemental Figure S1B-C). We hypothesized that *CD19*^low^ tumors may be heterogeneous, consisting of significant proportions of *CD19*⁻ tumor cells, whose elimination depends on Ag-independent CAR-T killing. To test this, we used knock-out (KO) cell lines to assess CAR-T on-target versus bystander killing in a reductionist system: αCD19 CAR-T were co-culture with CD19^+^ and CD19⁻ (1:1) human lymphoma cells, where CD19⁻ cells were either FAS^+^ or FAS⁻ (Figure 1F). On-target clearance of CD19^+^ cells (on-target killing) was observed, along with extensive clearance of CD19⁻ cells only when admixed with CD19^+^ cells (bystander killing), and in a highly Fas-dependent manner, supporting Ag-independent killing in mixed populations (Figures 1G-H). To determine if improved survival among patients with *FAS*^high^ lymphomas was specific to the CAR-T context, we similarly assessed outcomes of a separate cohort of patients receiving standard-of-care chemotherapy and therein observed no differences by *CD19* or *FAS* expression (Figures 1I-J). These data suggest that tumoral Fas signaling enhances therapeutic efficacy of CAR-T therapy in lymphoma, notably in those with heterogeneous Ag expression.

### CAR-T bystander killing prolongs survival in mice with Ag-heterogeneous tumors via Fas/FasL, antigen stimulation, and cell contact

To confirm the necessity of Fas signaling in CAR-T–mediated control of antigen-heterogeneous tumors, we transitioned to *in vivo* functional modeling. Mice were challenged either intravenously or subcutaneously with 80:20% mixtures of CD19^+^:CD19⁻ lymphoma cells, with cohorts receiving CD19⁻ bystander cells that were either Fas-sufficient (CD19⁻Fas^+^) or Fas-deficient (CD19⁻Fas⁻), prior to CAR-T transfer (Figure 2A). In the OCI-Ly8 tumor model, the median survival for 80:20% FAS⁻ tumor-bearing mice was 20.5 days with no therapy (n=6) and 21 days with CAR-T therapy (n=9), comparable to the survival of mice with 100% CD19^+^ tumors receiving no therapy (n=6, 21 days; Figure 2B). By contrast, 80:20% FAS^+^ tumor mice showed improved outcomes, with 22-day median survival with no therapy (n=8) versus 36 days with CAR-T therapy (n=8), approaching the survival of 100% CD19^+^ tumor mice receiving CAR-T therapy (n=6, 38.5 days; Figure 2B) and paralleling the patient data (Figures 1D-E). Similar results were observed in syngeneic A20 tumor model: no median survival difference was seen in the Fas^+^ and Fas⁻ mixed-tumor control groups receiving no CAR-T therapy (32 vs. 25 days; p=0.70), but median survival was significantly prolonged in Fas^+^ over Fas⁻ mice treated with CAR-T (45.5 vs. 37 days; p=0.03; n=7-8; Figure 2C).

**Figure 2.**
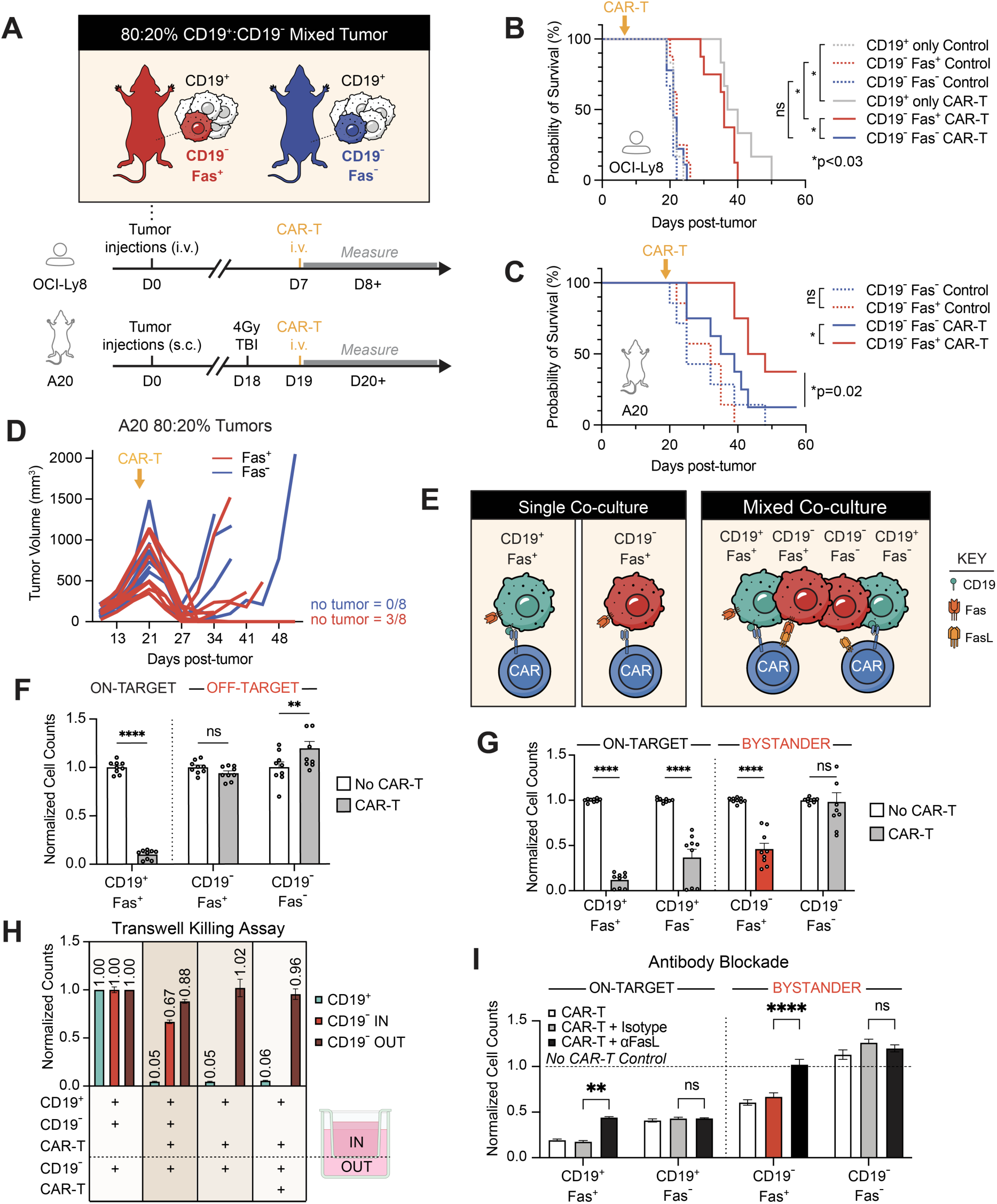
CAR-T bystander killing prolongs survival in mice bearing mixed-Ag tumors and is dependent on Ag-stimulation, Fas/FasL signaling, and cell contact. **(A)** Experimental design for mixed-tumor models with 80:20 CD19⁺:CD19⁻ tumors, using human (upper arrow) or mouse (lower arrow) systems, each containing either Fas⁺ (red) or Fas⁻ (blue) bystander populations. **(B)** Kaplan-Meier survival curves for OCI-Ly8 (human) and **(C)** A20 (mouse) bearing mixed-antigen tumors with either Fas⁺ (red) or Fas⁻ (blue) bystander populations, treated with or without CAR-T cells (n=6-9 mice per group). Statistical significance determined by Gehan-Breslow-Wilcoxon log-rank test; ns, not significant; *p<0.03. **(D)** Individual tumor growth curves for CAR-T treated mice bearing 80:20 mixed-antigen tumors. “No tumor” indicates the number of mice without palpable Fas⁻ (top) or Fas⁺ (bottom) tumors at the end of the experiment. **(E)** Left: Schematic of single-population co-cultures containing either CD19⁺ (green) or CD19⁻ (red) tumor cells with CAR-T (blue). Right: Schematic of mixed-population co-cultures containing equal proportions of CD19⁺Fas⁺, CD19⁺Fas⁻, CD19⁻Fas⁺, and CD19⁻Fas⁻. **(F-G)** Normalized cell counts for tumor populations following co-culture. **(F)** Single-population wells or **(G)** mixed-population wells, with bystander killing of CD19⁻ Fas⁺ cells highlighted in red. n=5 independent experiments. Dots indicate replicates; bars show mean ± s.e.m. Statistical significance determined by two-way ANOVA with Tukey correction. **(H)** Transwell co-culture assay. Conditions are grouped by whether CD19⁺ targets and CAR-T were located inside the same transwell compartment (IN) or separated from CD19⁻ bystander cells across the transwell membrane (OUT). See Supplemental Figure S2F for experimental schematic. **(I)** Antibody-blockade bystander killing assay. The dotted line indicates cell counts in the No CAR-T condition, and bystander killing of CD19⁻Fas⁺ cells is highlighted in red. n=3 independent experiments. Statistical significance determined by two-way ANOVA with Tukey correction. Dots indicate replicates; bars show mean ± s.e.m. Statistical significance was determined by log-rank test (B–C) and two-way ANOVA with multiple-comparisons correction as indicated (D–I); ns, not significant; *p < 0.03; **p < 0.002; ***p < 0.0002, ****p < 0.0001.

In the A20 model, on-target CD19^+^ cells were modified to lack CD20, enabling differentiation between “inborn” Ag-loss during therapy (CD19⁻CD20⁻) from the “engineered” parental CD19⁻ cells with intact CD20 (CD19⁻CD20^+^; Supplemental Figure S2A). After up to 3 weeks of tumor growth, untreated control mice maintained an average composition similar to initial composition (80:11% to 84:9%; n=8), signifying minimal growth advantage regardless of CD19, CD20, or Fas expression and consistent with survival patterns observed across R-CHOP treated patient cohorts (Supplemental Figure S2A, Figures 1I-J). Relapsing tumors following CAR-T therapy were overwhelmingly CD20^+^, indicating that the CD19⁻ tumor cells derive from selective pressure towards pre-existing CD19⁻ cells, as seen in recent reports,^41^ rather than mechanisms such as trogocytosis of target Ag (Supplemental Figure S2A). The impressive survival results spurred further evaluation of the mechanisms of CAR-T bystander killing.

Co-culturing αCD19 CAR-T effectors and up to four tumor cell populations: CD19^+^Fas^+^, CD19^+^Fas⁻, CD19⁻Fas^+^, and CD19⁻Fas⁻ (Figure 2E), we observed that when CAR-T effectors were co-cultured with single cancer cell populations, 90% of CD19^+^ targets were eliminated, with no killing of CD19⁻ bystander cells, confirming the lack of off-target killing (Figure 2F). In contrast, when all four target populations were co-cultured together with CAR-T effectors, 90% of CD19^+^Fas^+^ and 73% of CD19^+^Fas⁻ targets were killed, while 54% of CD19⁻Fas^+^ (red bar) and 2% of CD19⁻Fas⁻ bystander cells were cleared (Figure 2G, Supplemental Figure S2B). The remarkable efficiency of bystander killing against another syngeneic lymphoma line, EZB-P6, as well as the human OCI-Ly8 results, validated Fas-mediated CAR-T bystander killing across multiple tumor models (Supplemental Figures S2C-E). These data clearly demonstrate that bystander killing depends on proximity to nearby Ag^+^ cells and tumoral Fas expression.

Since multiple soluble factors can induce bystander killing—and certain forms of soluble FasL (sFasL) may induce killing^42^—we conducted transwell studies to understand whether direct cell-contact is required (Figure 2H, Supplemental Figure S2F). CAR-T cells effectively cleared 95% of CD19^+^Fas^+^ cells in the transwell setup (Figure 2H, 2^nd^ group) and induced bystander killing of 33% of CD19⁻Fas^+^ cells within the same compartment (Figure 2H, 2^nd^ group). However, CD19⁻Fas^+^ cells exposed to the same media, but separated by the transwell from CD19^+^Fas^+^ targets and CAR-T cells, exhibited minimal killing (12%, Figure 2H, 2^nd^ group), as did CD19⁻Fas^+^ cells exposed directly to CAR-T cells without CD19^+^Fas^+^ targets (4%; Figure 2H, 4^th^ group). These findings demonstrate that Ag-stimulation and cell-cell contact are required for effective CAR-T bystander killing.

To assess whether Fas receptor expression confers a cell-intrinsic growth advantage,^43^ αFasL blocking antibody was added to co-cultures and completely abrogated differences in killing between Fas^+^ and Fas⁻ cells (Figure 2I). Surprisingly, blocking antibodies against IFNγ, TNFα, and TRAIL-R had little effect on bystander killing (Supplemental Figure S2G). Collectively, these results demonstrated that CAR-T bystander killing requires Ag-stimulation, Fas-signaling, and direct cell contact.

### Rapid, contact-dependent bystander killing is mediated by Fas during a transient therapeutic window

To track the *in vivo* dynamics of CAR-T bystander killing in real-time, we engineered luciferase-expressing (Luc^+^) CD19⁻ tumor cells and implanted 80:20% CD19^+^:CD19⁻ mixed tumors with Fas⁻CD19⁻Luc^+^ and Fas^+^CD19⁻Luc^+^ components on opposite flanks of Balb/c Rag^-/-^ mice (Figure 3A). Dual-flank tumors allow direct, intra-subject comparison of bystander population fates during therapy accounting for inter-subject variability in CAR-T activity (Figures 3A-C). Therefore, each mouse served as its own control, with left-sided Fas⁻ CD19⁻ compared to right-sided Fas^+^ CD19⁻ Luc signal (Figure 3C). IVIS imaging dynamically visualized bystander killing over time (Figure 3B, Supplemental Figure S3A): prior to CAR-T therapy, CD19⁻Fas⁻ and CD19⁻Fas^+^ Luc signals were comparable; following CAR-T infusion, both tumors exhibited a marked decrease in Luc signal persisting for ∼10 days, indicating bystander killing (Figure 3C, upper panel). The real-time monitoring reveals what the non-Luc studies (Figure 2C) cannot: there is a significant component of Fas-independent bystander killing *in vivo* (Figure 3B, left tumors; Figure 3C, blue line) although insufficient to clear heterogeneous tumors and of lesser magnitude compared to Fas-dependent bystander killing (Figure 3B, right tumors; Figure 3C, red line). After the initial window of bystander killing, the Fas⁻ bystander signal re-emerged with significantly greater magnitude than the Fas^+^ signal (p=0.018) and persisted through day 28 post-CAR (p=0.016; Figure 3C, lower panel). While the mice exhibited macroscopic tumor regressions following CAR-T therapy and clear evidence of bystander killing, CD19⁻ tumor relapse ultimately contributed to their death (Supplemental Figures S3B-D). Whereas Fas-independent mechanisms contribute to early bystander tumor control *in vivo*, sustained bystander clearance *in vivo* appears more Fas-dependent.

**Figure 3.**
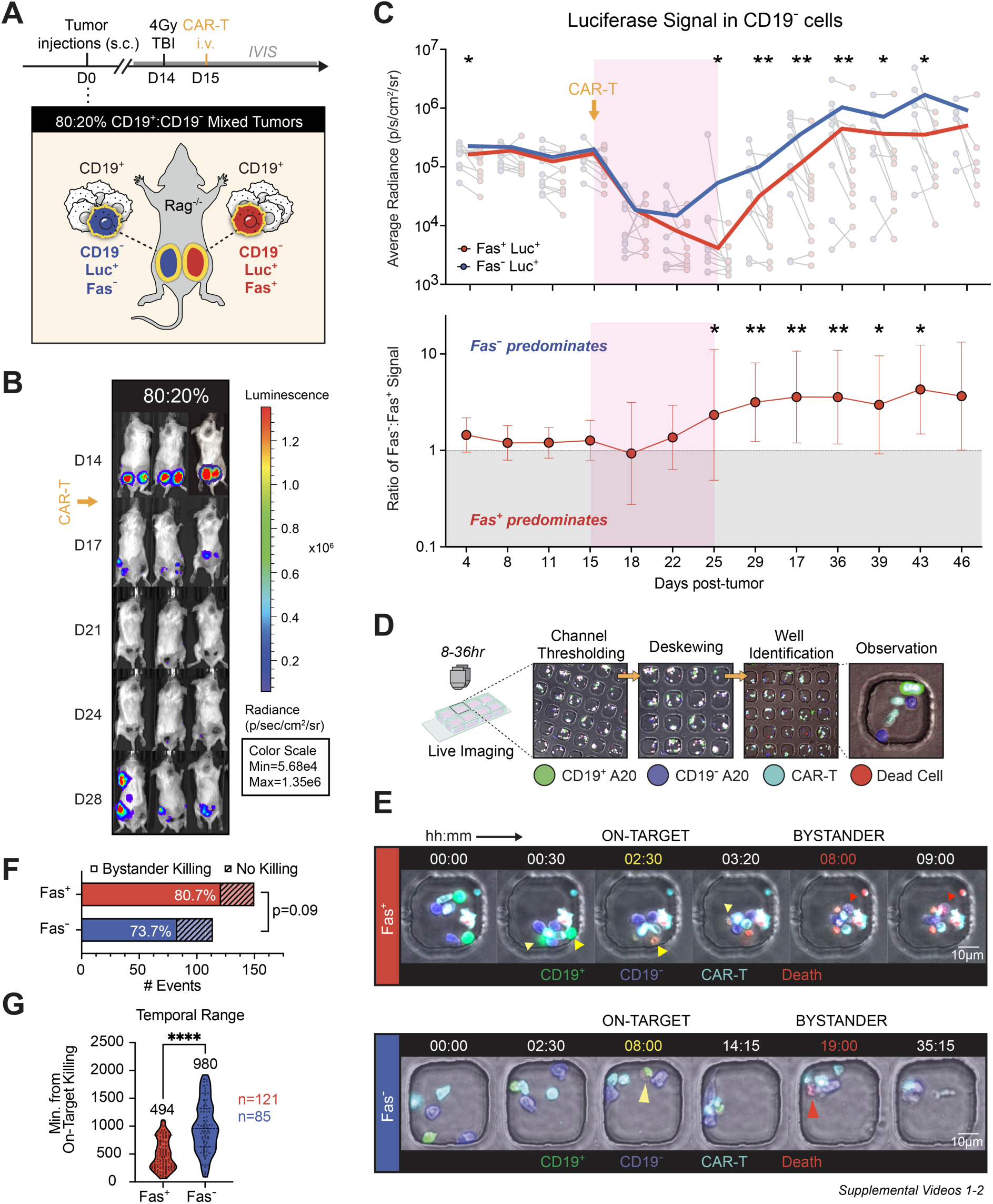
Spatiotemporal analysis of CAR-T bystander killing reveals a 10-day therapeutic window *in vivo* and accelerated death of Fas⁺ compared to Fas⁻ cells. **(A)** Schematic of the dual-flank mixed-tumor model. Rag⁻/⁻ mice were implanted with 80:20 CD19⁺:CD19⁻ luciferase⁺ (Luc^+^) tumors containing either Fas⁺ (yellow-outlined red) or Fas⁻ (yellow-outlined blue) CD19⁻ cells on opposite flanks. Mice received 4 Gy total body irradiation one day before intravenous CAR-T transfer, followed by longitudinal IVIS imaging to track the CD19⁻ bystander populations. **(B)** Representative IVIS images of mice bearing dual-flank tumors after CAR-T therapy, showing Fas^+^ (right flank) and Fas⁻ (left flank) CD19⁻Luc^+^ bystander populations over time. **(C)** Top: Average luciferase signal from Fas⁻ and Fas⁺ CD19⁻Luc^+^ bystander populations (foreground), with paired per-mouse signals shown in the background. Bottom: Ratio of Fas⁻ to Fas⁺ tumor signal over time; values >1 indicate predominance of Fas⁻ tumors. Therapeutic window of bystander killing denoted by shaded red box. **(D)** Schematic of PDMS microwell live-imaging assay used to monitor cell–cell interactions at single-cell resolution. Representative still images show CD19^+^ tumor cells (green), CD19⁻ tumor cells (blue), CAR-T (cyan), and dead cells (red). **(E)** Representative still images from live imaging of co-cultures containing CAR-T cells with either Fas⁺ or Fas⁻ bystander cells. Yellow arrows mark CAR-T–mediated on-target killing; red arrows mark bystander killing. Time stamps are shown as hh:mm. Scale bar, 10µm (applies to all images). **(F)** Frequency of bystander killing events per well-of-interest, defined as wells containing CAR-T, CD19⁺, and CD19⁻ cells. Statistical significance was assessed by Welch’s two-tailed t-test; n=3 independent experiments. **(G)** Violin plot showing the kinetics of bystander killing in Fas⁺ versus Fas⁻ co-cultures, measured as the interval between on-target killing and subsequent bystander cell death. Statistical significance was assessed by unpaired two-tailed t-test. *p < 0.03, **p < 0.002, ***p < 0.0002, ****p < 0.0001.

The surprising temporal dynamics of bystander killing Fas-dependence *in vivo* prompted deeper analysis of the kinetics of CAR-T bystander killing at single-cell resolution, using live imaging in microwell chambers (Figure 3D).^44^ CAR-T cells (GFP- or dye-labeled) were co-cultured with a mixture of dye-labeled CD19^+^ targets and CD19⁻ bystander cells, with 3-6 cells loaded per microwell (Figure 3D). Cell death was identified by propidium-iodide influx and/or membrane blebbing (Figure 3D). Bystander killing events were defined as death of a CD19⁻ cell following CAR-T-mediated killing of a CD19^+^ cell. We hypothesized that CD19⁻Fas^+^ cells would be killed more frequently and with faster kinetics than Fas⁻ counterparts; we observed a trend of more frequent bystander killing against Fas^+^ compared to Fas⁻ cells (81% vs. 73%, p=0.09, one-sided Fisher’s exact test; Figure 3F). However, Fas^+^ cells were killed with markedly greater rapidity: on average 494min (8.2 h) after on-target killing, with killing occurring as early as 3 hours and up to 20 hours from on-target killing, while Fas⁻ cells were killed on average 980min (16.3 h) and up to 1845min (30.75 h) after on-target killing (Welch’s t-test, p<0.0001; Figure 3E, Supplemental Videos S1-2). Assessing the spatial scope of bystander killing events, we observed that many such events occurred in the context of CAR-T:CD19^+^:CD19⁻ “triads” (Supplemental Figure S3E), a feature previously noted by Liadi and colleagues.^45^ Quantifying this, CAR-T:CD19^+^:CD19⁻ triads were observed in 56% of Fas^+^ wells but only 27% of Fas⁻ wells (p<0.0001; Supplemental Figure S3F). These findings suggest that Fas-dependent bystander killing occurs more rapidly and may preferentially rely on sustained cell-contact.

As extrinsic apoptotic signaling can result in apoptosis within one hour of receptor engagement,^46^ we posit that the faster kinetics of Fas-mediated CAR-T bystander killing may be attributed to the cell-contact induction of apoptosis reliant upon activation-induced FasL expression.^47–50^ Notably, FasL upregulation on CAR-T required the presence of CD19^+^ target cells and was expressed on the surface for up to 12 hours after engagement (Supplemental Figure S3G-H), which would permit cell death-inducing FasL-Fas multimerization.^51,52^ By contrast, the delayed killing of Fas⁻ bystanders suggests involvement of slower mechanisms (e.g. cytokines).^53^ Together, these studies support a model in which Fas-mediated bystander killing *in vitro* and *in vivo* is faster, localized, and contact-dependent compared to Fas-independent mechanisms.

### Potentiating Fas signaling or FasL expression enhances bystander killing in tumor cells, which can be further improved by decoupling Fas signaling in CAR-T cells

Given the clear contribution of Fas-FasL signaling to CAR-T bystander killing, we hypothesized that enhancing Fas signaling—either by blocking downstream negative regulators or by promoting FasL stabilization—could prevent the outgrowth of CD19⁻ tumors that contributed to death (Supplemental Figures S3B-C). The inhibitor of apoptosis (IAP) protein family emerged as a lead target due to its frequent overexpression in lymphoma and evidence of improved CAR-T efficacy with IAP inhibition (IAPi; Figure 4A, Supplemental Figure S4A-B).^32,34^ Treatment with increasing doses of birinapant, a clinically studied IAPi, increased apoptosis markers (cleaved caspase-3 and Annexin V) from 46% to 56% in CD19⁻Fas^+^ lymphoma cells admixed with CD19^+^ cells and CAR-T cells, but minimally affected CD19⁻Fas⁻ cells or CD19⁻ cells without CAR-T co-culture (Figure 4B). Correspondingly, the frequency of CD19⁻Fas^+^ cells was IAPi-dose-dependently reduced from 44% to 16%, with no change in CD19⁻Fas⁻ cells (Figure 4C). Similar effects were noted with other IAPi (CUDC-427 and xevinapant) and in OCI-Ly8 human cells (Supplemental Figures S4D-I). Synergy analysis confirmed a significantly greater-than-additive effect of IAPi on CAR-T bystander killing (Figure 4D, Bliss synergy score: 10.394).

**Figure 4.**
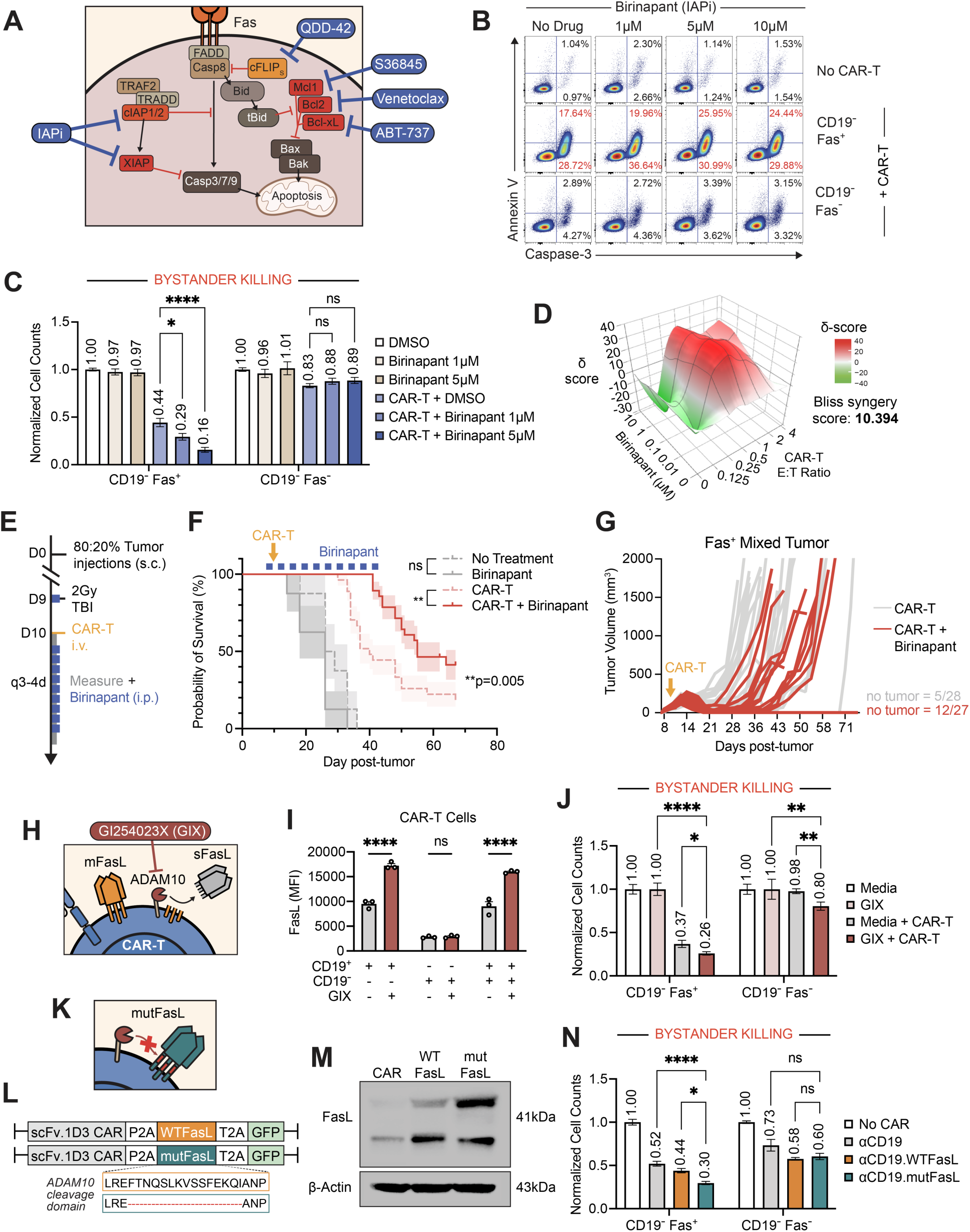
IAP inhibition and FasL stabilization potentiate Fas signaling to enhance CAR-T bystander killing. **(A)** Schematic of the Fas signaling pathway highlighting key regulatory proteins and corresponding pharmacologic inhibitors. **(B)** Representative flow cytometry plots showing cleaved caspase-3 and Annexin V staining of CD19⁻Fas⁺ and CD19⁻Fas⁻ bystander tumor cells co-cultured with CAR-T cells in the presence or absence of birinapant (Bir). **(C)** Normalized viable cell counts of CD19⁻Fas⁺ and CD19⁻Fas⁻ tumor cells in mixed co-culture with CAR-T across increasing birinapant doses. *n* = 5 independent experiments. **(D)** Synergy analysis using Bliss scoring. The x-axis shows increasing effector-to-target (E:T) ratios, the y-axis shows birinapant doses, and the z-axis shows delta-scores. A Bliss synergy score >10 indicates a greater-than-additive effect. **(E)** Schematic of the experimental design to assess birinapant potentiation of CAR-T bystander killing in mixed-antigen tumors. **(F)** Survival curves for mice bearing 80:20 Fas⁺ mixed tumors treated with or without CAR-T in the presence or absence of birinapant. Mice received 2 Gy TBI on day 9 and CAR-T transfer on day 10; birinapant (15 mg/kg) was dosed twice weekly for 10 doses starting on day 9. No CAR-T cohorts, n=8 per group; CAR-T cohorts, n=28 (no birinapant) and n=27 (birinapant). Statistical significance was determined by log-rank test; 95% CI shown by shaded area above and below survival line. **(G)** Individual tumor growth curves for mice in (F). “No tumor” indicates the number of mice without palpable tumors at the end of the experiment, shown separately for no birinapant (top) or birinapant-treated (bottom) groups. **(H)** Schematic of FasL shedding by ADAM10 and its inhibition by GI254023X (GIX). **(I)** Surface FasL expression on CAR-T cells with or without GIX treatment in the presence of CD19^+^, CD19⁻, or mixed target cell populations. **(J)** Normalized viable cell counts of CD19⁻Fas⁺ and CD19⁻Fas⁻ tumor cells in mixed co-culture with CAR-T across increasing GIX doses. *n* = 3 independent experiments. **(K)** Schematic of non-cleavable mutFasL resisting ADAM10-mediated shedding. **(L)** Schematic of retroviral vectors for αCD19 CAR constructs overexpressing either wild-type FasL (WTFasL, orange) or non-cleavable mutant FasL (mutFasL, teal). The amino acid sequence of the ADAM10 cleavage domain is shown below the constructs. **(M)** FasL protein expression in CAR-T cells. mutFasL migrates at slightly higher molecular weight due to non-cleavable modification. **(N)** Normalized viable cell counts of CD19⁻Fas⁺ and CD19⁻Fas⁻ tumor cells in mixed co-culture with WTFasL CAR-T versus mutFasL CAR-T. *n* = 3 independent experiments. Dots represent technical replicates, bars indicate mean ± s.e.m. Statistical significance determined by two-way ANOVA with Tukey’s multiple comparisons. *p < 0.03, **p < 0.002, ***p < 0.0002, ****p < 0.0001.

IAPi effects on *in vivo* CAR-T bystander killing have been previously studied in xenograft glioblastoma models in immunodeficient recipients, though these are confounded by xenograft-versus-host effects.^54^ As IAPi effects on *in vivo* CAR-T bystander killing have not been studied in syngeneic, immunocompetent recipients, we assessed this in the mixed-tumor model (Figure 4E). Mice were challenged with 80:20% CD19^+^:CD19⁻ mixed tumors and treated with either no CAR-T or 7.5 x 10^5^ CAR-T cells, with or without birinapant (15 mg/kg, i.p.) administered twice weekly for 10 doses. In the no CAR-T cohorts, median survival was similar with or without birinapant (26 vs. 27.5 days, p=0.115; n=8; Figures 4E-F, Supplemental Figure S4C). In CAR-T-treated mice, median survival was markedly better with versus without birinapant (55 vs. 44 days, p=0.005, n=27, n=28; Figure 4F). Impressively, the addition of birinapant to CAR-T therapy delayed macroscopic tumor growth and even led to complete tumor regressions in 12/27 mice as compared to 5/28 without IAPi (Figure 4G). As noted, nearly all tumoral relapse in these 80:20 mixed-Ag tumors were due to escape by CD19⁻ tumor cells (Supplemental Figure S2A), implicating IAPi as effective potentiators of CAR-T-mediated bystander killing.

As an orthogonal approach to increasing Fas receptor aggregation and signaling, we assessed the effects of increasing CAR-T membrane-bound FasL (mFasL) expression. Since the trimeric form of mFasL induces target-cell cytotoxicity more effectively than the soluble/cleaved form (sFasL),^52,55^ we hypothesized that preventing mFasL cleavage may increase bystander killing. Treatment with GI254023X (GIX), an inhibitor that blocks mFasL shedding by ADAM10 metalloproteinase (Figure 4H),^56^ significantly increased mFasL expression on Ag-stimulated CAR-T cells with corresponding decrease in sFasL (Figures 4I-J, Supplemental Figure S4J). The addition of GIX to co-culture led to an 11% increase in bystander killing (Figure 4K), but also increased Fas-independent killing—perhaps due to off-target effects on other ADAM10 substrates (e.g., CD44, PD-L1, and MICA/B).^57^ To directly test mFasL stabilization, we constructed a CAR-T cell with constitutive expression of wild-type mFasL (WTFasL) and non-cleavable mFasL^58^ (mutFasL; Figures 4K-L). The expression of the WTFasL and mutFasL was confirmed by western blot (Figure 4M). Compared to αCD19 CAR-T (48% bystander killing), αCD19.WTFasL and αCD19.mutFasL CAR-T increased killing to 56% and 70%, respectively (Figure 4N). Although some Fas-independent bystander killing was also notable in these assays, it was not significantly increased over αCD19 CAR-T (Figure 4N), and may reflect FasL-mediated CAR-T activation and greater cytokine release, which warrants further study.^59,60^

To determine whether bystander killing is exclusive to CAR-T-based systems, we tested bystander killing with BsAb-redirected T cells (Figure 5A). In the OCI-Ly8 model, we confirmed that CD20^+^ and CD20⁻ cells were insensitive to off-target killing by T cell effectors, but sensitized to both on-target killing and bystander killing with Epcoritamab, a CD20xCD3 BsAb (Figure 5B). As observed with CAR-T bystander killing, the OCI-Ly8 CD20⁻ Fas⁻ cells were more resistant than their CD20⁻ Fas^+^ counterparts to BsAb bystander killing (Figure 5B). Because Fas-susceptibility varies across human lymphoma lines,^61^ we next tested Raji cells, which are significantly more Fas-resistant (Figures 5C-E). In this setting, the addition of Epcoritamab alone did not elicit appreciable bystander killing (Figure 5F). Bystander killing only emerged when combined with birinapant, highlighting that tumoral Fas sensitization with IAPi is crucial to initiating bystander killing in resistant tumor contexts (Figure 5F).

**Figure 5.**
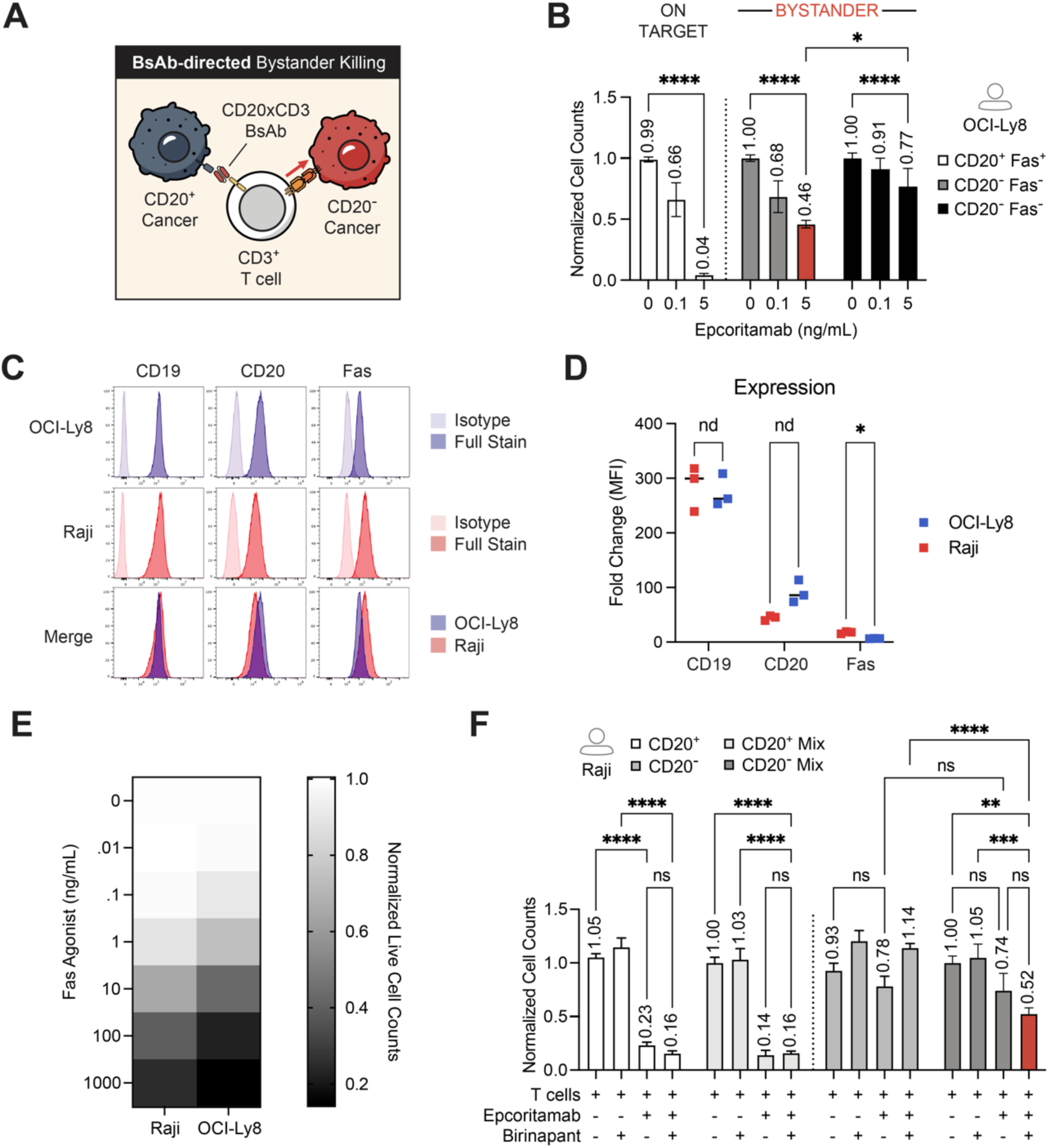
Bispecific-redirected T cells mediate bystander killing of antigen-negative cells, which may require Fas sensitization with IAP inhibition. **(A)** Schematic of a CD3^+^ T cell engaging a CD20^+^ tumor cell through a CD20xCD3 bispecific antibody (BsAb) leading to Fas-mediated bystander killing on an adjacent CD20-cell. **(B)** Normalized viable cell counts of CD20^+^, CD20⁻Fas^+^, and CD20⁻Fas⁻ OCI-Ly8 human lymphoma cells following co-culture with increasing doses of CD20xCD3 BsAb Epcoritamab. Bystander killing of CD20⁻Fas^+^ cells is highlighted in red. Multiple t-tests were conducted on pre-specified comparisons, without correction. **(C)** Surface expression of CD19, CD20, and Fas in two human lymphoma cell lines OCI-Ly8 and Raji. **(D)** Mean fluorescence intensity (MFI) fold-change of surface expression in **(C)**. **(E)** Heatmap of normalized viable cell counts of OCI-Ly8 and Raji cells following 72 h co-culture with αFas agonist antibody. Boxes represent mean values. n=3 technical replicates. **(F)** Normalized viable cell counts of CD20^+^ and CD20⁻ Raji cells after 48 h co-culture with Epcoritamab in the presence or absence of birinapant. Bystander killing of CD20⁻ cells is highlighted in red. Dots indicate replicates; bars show mean ± s.e.m. Statistical significance was determined by two-way ANOVA with correction for multiple comparisons. *p < 0.05, **p < 0.01, ***p < 0.001, ****p < 0.0001.

Since IAP also regulate T cell survival,^62,63^ we assessed potential extra-tumoral effects of IAPi. Following Ag-stimulation, Fas expression was upregulated to a greater degree on CD4^+^ than CD8^+^ CAR-T (Figures 6A-B). Surprisingly, we also observed that CD4^+^ much more so than CD8^+^ CAR-T expressed greater levels of cleaved-capase-3 (Figure 6C) corresponding to decreased cell numbers with birinapant co-treatment (Figure 6D), suggestive of Fas-mediated fratricide (Figure 6E), as seen with CD28-based CAR-T and synthetic TCR T models.^64,65^ Despite CD4^+^ CAR-T loss with IAPi, overall bystander killing increased (Figure 4C). Moreover, as CD4^+^ CAR-T alone were able to induce bystander killing in addition to CD8^+^ CAR-T (Supplemental Figure S5A), we asked whether decoupling Fas signaling in CAR-T could preserve CD4^+^ viability and thereby further improve the efficacy of IAPi.^66,67^ Indeed, Fas-knockout (FasKO) CAR-T showed restored CD4^+^ counts (Figures 6E-G, Supplemental Figure S5B) and significantly superior bystander killing in the setting of IAPi compared to non-KO CAR-T cells (p<0.0001; Figure 6H). Accordingly, the most pronounced effects were observed with FasKO in combination with IAPi (Figure 6H) and αCD19.mutFasL CAR (Figure 6I), increasing clearance of bystander populations over 80%. In sum, these findings establish that enhanced Fas signaling in tumor cells via IAPi or FasL stabilization, and decoupling Fas sensitivity in T cells synergistically boosts CAR-T bystander killing.

**Figure 6.**
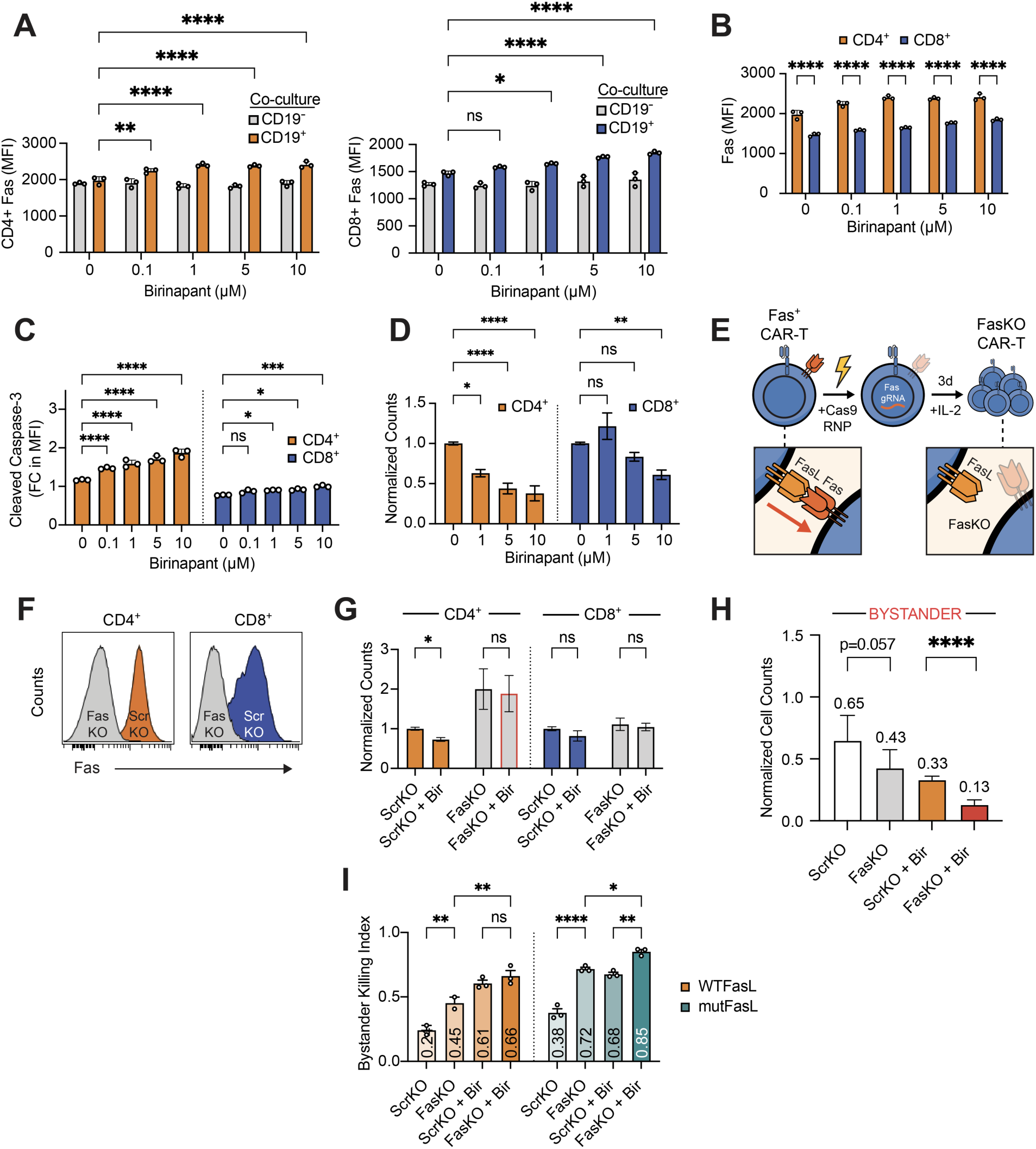
IAP inhibition enhances bystander killing but induces CD4⁺ T cell fratricide, which can be rescued by Fas knockout. **(A)** Viable CD4⁺ and CD8⁺ CAR-T counts after 48 h co-culture with CD19⁺ or CD19⁻ target cells in the presence of increasing doses of birinapant. **(B)** Surface Fas expression on CD4⁺ and CD8⁺ CAR-T after birinapant treatment. **(C)** Percentage of cleaved caspase-3⁺ CD4⁺ and CD8⁺ CAR-T after co-culture ± birinapant. Increased apoptosis is observed in CD4⁺ cells. **(D)** Viable cell counts of CD4⁺ and CD8⁺ CAR-T after co-culture with increasing doses of birinapant. **(E)** Schematic of the mechanism of CD4⁺ T cell fratricide via Fas-FasL and the Fas knockout (FasKO) strategy to preserve T cell viability. **(F)** Surface Fas expression on scramble gRNA control (ScrKO) versus FasKO CAR-T. **(G)** Viable CD4⁺ and CD8⁺ CAR-T counts after co-culture with ScrKO or FasKO CAR-T in the presence or absence of birinapant. CD4^+^ CAR-T FasKO with birinapant is highlighted in red. *n* = 2 independent experiments. **(H)** Normalized viable cell counts of CD19⁻Fas^+^ tumor cells following 48 h co-culture with ScrKO or FasKO CAR-T in the presence or absence of birinapant (5 µM). *n* = 2 independent experiments. **(I)** Bystander killing index of CAR-T overexpressing wild-type Fasl (WTFasL) or non-cleavable mutant FasL (mutFasL), with or without FasKO, in the presence or absence of birinapant (5 µM). Dots indicate technical replicates; bars show mean ± s.e.m. Statistical significance was determined by two-way ANOVA with Tukey’s multiple comparisons (A-D, I) or by pre-specified t-test comparisons (G-H). *p < 0.05, **p < 0.01, ***p < 0.001, ****p < 0.0001.

### Bystander killing can be harnessed for non-lymphoma tumors by targeting tumor microenvironment antigens

Conceptually, CAR-T bystander killing requires local Ag-stimulation to eliminate adjacent Ag⁻ cancer tissue. This process is not limited to tumor-associated Ag (TAA) on tumor *cells*, but may also be triggered by tumor-microenvironment surface Ag (TME-Ag) on tumor-*adjacent* cells (Figure 7A).^68^ Targeting TME-Ag is an emerging immunotherapeutic strategy that can boost secondary, cancer-targeting CAR-T responses, as well as reshape the endogenous immune population in resistant tumors^69,70^ (Mateus-Tique et al., under review). However, contrary to findings that tumor-associated macrophage (TAM) clearance alone does not reduce tumor burden, utilizing αFOLR2 CAR-T^69^ to target FOLR2^+^ Raw264.7 macrophage cells alongside bystander FOLR2⁻ A20 cells resulted in remarkable bystander killing, which significantly improved with birinapant co-treatment (Figure 7B).

**Figure 7.**
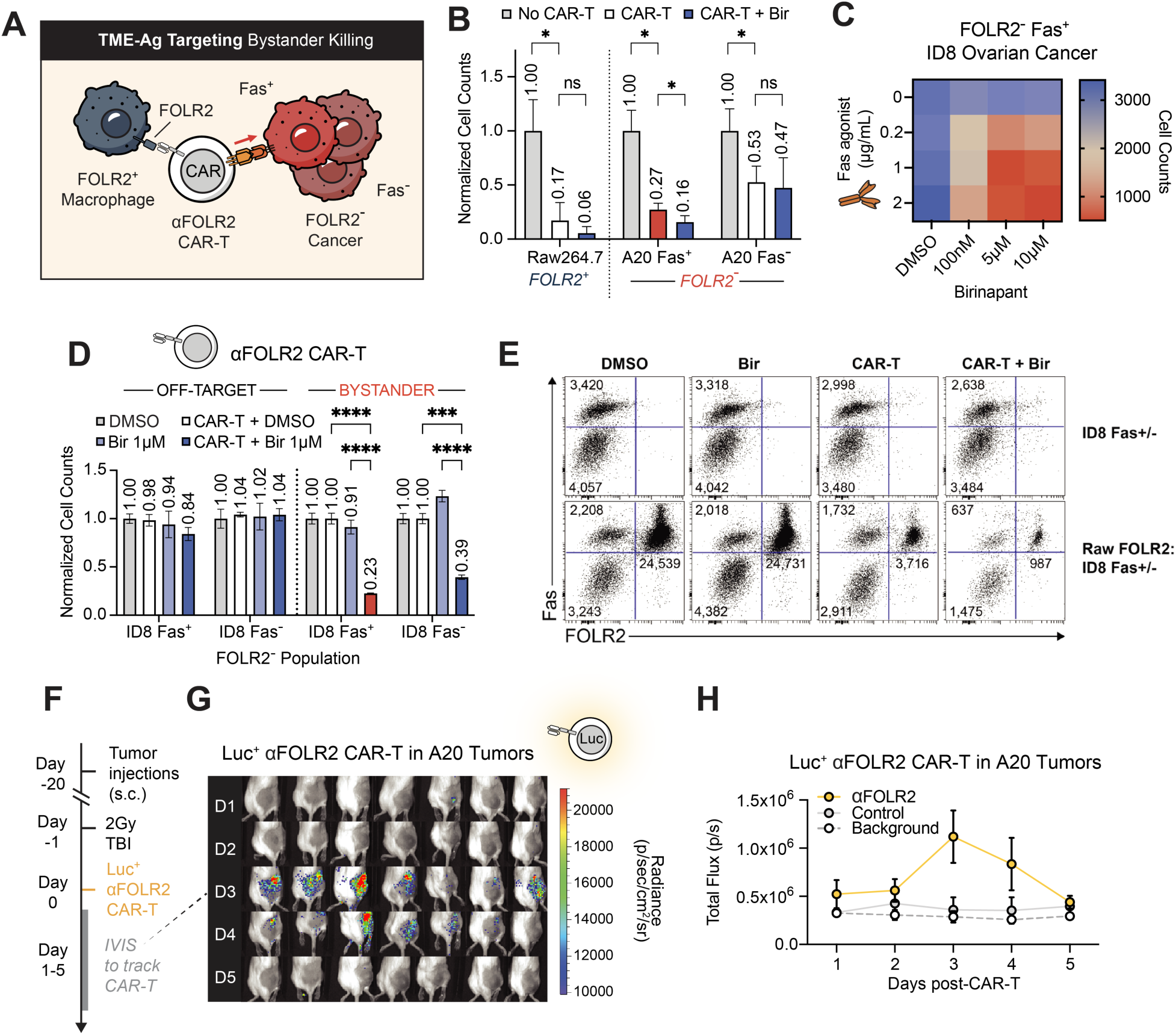
CAR-T targeting tumor-microenvironment antigens mediates bystander killing of tumor cells lacking uniform surface antigens and extend the platform to solid tumors. **(A)** Schematic of tumor microenvironment antigen (TME-Ag)-directed bystander killing. Anti-FOLR2 CAR-T cells target FOLR2⁺ tumor-associated macrophages (TAMs), leading to Fas-dependent killing of adjacent FOLR2⁻ tumor cells. **(B)** Normalized viable cell counts of FOLR2⁺ Raw264.7 macrophages and FOLR2⁻ A20 tumor cells after 48 h co-culture with anti-FOLR2 CAR-T in the presence or absence of birinapant (5 µM). Bystander killing of FOLR2⁻Fas^+^ A20 cells is highlighted in red. *n* = 2 independent experiments. **(C)** Heatmap of ID8 ovarian cancer cell viability after treatment with increasing doses of anti-Fas agonist, birinapant, or both. **(D)** Normalized viable cell counts of FOLR2⁺ Raw264.7 macrophages and FOLR2⁻ ID8 tumor cells after 48 h co-culture with anti-FOLR2 CAR-T in the presence or absence of birinapant (1 µM). Bystander killing of FOLR2⁻Fas^+^ ID8 cells is highlighted in red. *n* = 3 technical replicates. **(E)** Representative flow cytometry dot plots from the co-culture experiment in (D). **(F)** Experimental timeline for tracking Luc^+^ anti-FOLR2 CAR-T in mice bearing FOLR2⁻ A20 lymphoma tumors. **(G)** IVIS images of Luc^+^ anti-FOLR2 CAR-T localizing to FOLR2⁻ A20 lymphoma tumor. **(H)** Quantification of Luc^+^ anti-FOLR2 CAR-T from (G). Dots indicate replicates; bars show mean ± s.e.m. Statistical significance was determined by two-way ANOVA with Tukey’s multiple-comparisons test. *p < 0.05, **p < 0.01, ***p < 0.001, ****p < 0.0001.

Because TAM-targeting strategies are primarily applied to non-lymphoma solid tumors enriched in FOLR2^+^ TAM,^71^ we evaluated ID8 ovarian cancer cells. ID8 cells were resistant to Fas agonist antibody or birinapant treatment alone, but showed marked dose-dependent sensitivity to combination therapy (Figure 7C). Correspondingly, in co-culture experiments of FOLR2^+^ Raw264.7 cells with FOLR2⁻ ID8 cells, bystander killing was minimal without birinapant, and remarkable in its presence: 77% of FOLR2⁻ Fas^+^ and 61% of FOLR2⁻ Fas⁻ ID8 cells were bystander killed (Figures 7D-E). This effect may be driven by birinapant’s modulation of both apoptotic and NFκB pathways,^72^ especially given the observed Fas upregulation by IFNγ and TNFα (Supplemental Figure S6A). B16 melanoma cells were also susceptible to bystander killing when mixed with FOLR2^+^ Raw264.7 cells (Supplemental Figure S6B). Preliminary studies with a novel mouse BsAb further demonstrate the feasibility of targeting TME-Ag with CD3^+^ T cells and inducing bystander killing of A20 tumor cells *in vitro* (Supplemental Figures S6C-D). Validating this TAM-targeting approach *in vivo*, Luc^+^αFOLR2 CAR-T cells trafficked to FOLR2⁻ A20 tumors within three days of transfer, confirming effective tumor localization and proliferation (Figures 7E-F). These findings demonstrate that bystander killing can also be triggered through TME-targeting strategies, establishing bystander killing as a generalizable mechanism applicable to non-lymphoma tumors lacking actionable TAA.

## Discussion

Ag escape remains a formidable challenge to durable CAR-T responses and is expected to become increasingly prevalent as TRT expand in clinical use.^73^ While strategies like multi-Ag targeting and next-generation CAR platforms seek to address Ag escape once it has occurred, the observation of durable responses in Ag-heterogeneous tumors suggests mechanisms already mitigating Ag escape. Here, we define Fas-mediated CAR-T bystander killing as a potent, tunable mechanism that enables cytotoxicity against Ag-heterogeneous tumors *a priori*.

We demonstrate that *FAS* expression is a key determinant of survival in CAR-T-treated DLBCL patients with low *CD19* expression, but not in those on standard chemotherapy, suggesting a specific contribution of Fas-signaling in Ag-independent clearance by CAR-T therapy. Using mouse and human models, we confirm that CAR-T bystander killing is dependent on Ag-stimulation, tumoral Fas expression, and direct cell contact. Furthermore, spatiotemporal tracking by microwell imaging revealed faster and more frequent killing kinetics through Fas-dependent over Fas-independent pathways.

Compared to Fas-mediated triad-based killing, Fas-independent mechanisms may rely on diffusion or the contribution of innate immune cell signaling and thereby be less spatially and temporally efficient.^22^ This effect was recapitulated *in vivo*, where Fas^+^ bystander tumors show prolonged tumor suppression compared to Fas⁻ counterparts. Together, these data support a model where multiple signals contribute to bystander killing, but Fas-FasL signaling is dominant and temporally constrained.

Critically, we show that the Fas-signaling axis can be pharmacologically and genetically enhanced. IAPi such as birinapant sensitize Fas^+^ tumor cells to CAR-T bystander killing without direct drug toxicity, leading to improved clearance and extended survival *in vivo*. We further demonstrate BsAb-directed bystander killing, generalizing the effect to multiple T cell modalities. Since IAP are also expressed in CAR-T cells, an increase in CD4^+^ T cell fratricide was also observed, even in the setting of improved bystander clearance. This motivated the decoupling of Fas-signaling in CAR-T cells by FasKO, which rescued T cells from fratricide and further improved the IAPi synergistic effect. Further enhancement is achieved by increasing mFasL expression on T cells, either by preventing ADAM10 cleavage with small molecule inhibitor or expression of non-cleavable, membrane-bound FasL. The combination of mFasL stabilization and T cell FasKO yields significantly increased clearance of Ag-tumor populations *in vitro*.

To our knowledge, this is also the first report of CAR-T bystander tumor killing via targeting of non-tumor cell Ag. αFOLR2 CAR-T directed against FOLR2^+^ TAMs elicit clearance of adjacent, FOLR2⁻ tumor cells, an effect potentiated by IAPi and observed in ovarian cancer models. This proof-of-concept has two-fold benefit: the specific clearance of immunosuppressive cells and the bystander clearance of tumors that may suffer from the lack of universally-expressed Ag.

These findings have important implications: first, they strongly suggest that CAR-T efficacy in Ag-heterogeneous tumors may already rely, in part, on Fas-dependent bystander killing. Second, they establish that this mechanism is therapeutically actionable through pharmacologic sensitization and synthetic biology. Third, they demonstrate that targeted T cell therapies can be unleashed in historically Ag-deficient tumor settings by targeting tumor-extrinsic features and with T cell-redirecting activation.

Nonetheless, the durability of bystander killing appears temporally limited, perhaps due to the transient expression of mFasL and the contraction of CAR-T effectors over time.^74^ Moreover, while Fas-FasL signaling serves a dominant role in CAR-T bystander killing, Fas-independent mechanisms likely work in parallel and may be particularly relevant in solid tumor contexts where limited cell-cell contact restricts cytotoxicity. For example, diffusible mediators, such as IFNγ or TNFα, provide complementary modes of bystander killing; future work should investigate how Fas-dependent and -independent pathways interact, and whether combinatorial strategies can potentiate both contact-dependent and diffusible killing.

Beyond pathway synergy, anti-tumor T cells with dual-costimulatory CD28 and 4-1BB domains have shown efficacy in preventing Ag escape,^75,76^ possibly because of a difference in FasL dynamics.^77^ Integrative logic-gated platforms could also play a crucial role in extending bystander killing.^78,79^ For example, CAR-T cells could target a lowly-expressed TAA (e.g., EGFRvIII) and subsequently secrete BsAb against TME-A (e.g., FOLR2), enabling dynamic redirection of cytotoxicity and prolonging duration of bystander killing as primary Ag decreases and secondary Ag increases. Engineering dominant-negative Fas receptors^74^ or switch receptors^80^ may similarly rewire suppressive signals into activation cues that amplify effector function. Finally, *in vivo* programming of CAR-T cells or tumor-infiltrating lymphocytes to express TCR-driven stabilized FasL—or employing OFF-switch constructs^81^ to temporally regulate FasL expression—could offer more precise control over bystander killing to further reduce off-target toxicity in both hematologic and solid tumor settings.

Overall, our findings establish Fas-mediated bystander killing as a mechanistically-distinct and therapeutically-actionable axis of targeted T cell therapy. By shifting the focus from Ag-restriction to tumor Ag-proximity and tumoral death-signal tuning, this work provides a therapeutic framework to mitigate Ag escape. As CAR-T and other TRT continue to proliferate, leveraging bystander killing through rational Fas pathway modulation offers a compelling strategy to improve therapeutic durability, mitigate Ag escape, and enable targeting in otherwise Ag-intractable disease.

---

**Supplemental Figure S1. Correlation of death receptor signature (DRS) genes and survival outcomes in *CD19*^low^*FAS*^low^ DLBCL patients. Related to Figure 1.**

(A) Pairwise correlation heatmap of genes in the Death Receptor Signature (DRS).

(B) Event-free survival and (C) overall survival of DLBCL patients receiving CAR-T therapy stratified by low CD19 and Fas expression. Statistical significance determined by log-rank tests.

**Supplemental Figure S2. Fas signaling is necessary and sufficient for CAR-T bystander killing across human and murine systems. Related to Figure 2.**

**(A)** Representative flow cytometry plots of 80:20% mixed-antigen tumors before (left) and after (right) CAR-T treatment from the same experiment as in Figure 2C. Histogram shows surface Fas expression on CD19⁻CD20^+^ tumor cells following CAR-T therapy.
**(B)** Representative changes in tumor cell populations following mixed-population co-culture with CAR-T, as in Figure 2G.
**(C)** Normalized viable cell counts of CD19^+^Fas^+^, CD19⁻Fas^+^, and CD19⁻Fas⁻ EZB P6 tumor cells following mixed-population co-culture. n=2 independent experiments.
**(D)** Normalized viable cell counts of CD19^+^Fas^+^, CD19⁻Fas^+^, and CD19⁻Fas⁻ EZB P6 tumor cells following single-population co-culture. n=2 independent experiments.
**(E)** Normalized viable cell counts of CD19^+^Fas^+^, CD19⁻Fas^+^, and CD19⁻Fas⁻ OCI-Ly8 tumor cells following single-population co-culture with CAR-T. *n*=2 independent experiments.
**(F)** Schematic of the transwell assay (as in Figure 2H), with the corresponding type of killing observed in each condition labeled above and below the transwell.
**(G)** Normalized viable cell counts of CD19^+^Fas^+^, CD19^+^Fas⁻, CD19⁻Fas^+^, and CD19⁻Fas⁻ following mixed-population co-culture with CAR-T in the presence of various blocking antibodies. Statistical significance was determined by two-way ANOVA with Dunnett correction; *n*=2 independent experiments.

Dots indicate replicates; bars show mean ± s.e.m. Statistical significance was determined by two-way ANOVA with correction for multiple comparisons. *p < 0.03, **p < 0.02, ***p < 0.0002, ****p < 0.0001.

**Supplemental Figure S3. Real-time imaging of CAR-T bystander killing in vivo and in vitro reveals both Fas-dependent and -independent mechanisms. Related to Figure 3.**

**(A)** Extended IVIS images of mice from Figure 3B. Days post-tumor injection are labeled on the left.

**(B)** Individual tumor growth curves for Fas⁻ and Fas^+^ mixed-antigen tumors for mice in experiment shown in Figure 3A-C.

**(C-D)** Flow cytometry plots of pre-treatment (C) and post-treatment (D) mixed-antigen tumors showing GFP and CD19 expression. Fas+ tumors from the right flank of the mice are shown in the top row; Fas-tumors from the left flank of the mice are shown in the bottom; dLN, draining lymph node; aLN, axillary lymph node; IVIS images of mice are shown in the top right corner.

**(E)** Representative still images of CAR-T:CD19^+^:CD19⁻ triads from live imaging of co-cultures containing CAR-T cells with either Fas⁺ or Fas⁻ bystander cells, as in Figure 3E. Yellow arrows mark CAR-T–mediated on-target killing; red arrows mark bystander killing. Time stamps are shown as hh:mm. Scale bar, 10µm (applies to all images).

**(F)** Frequency of triad bystander killing events per well-of-interest, defined as wells containing CAR-T, CD19⁺, and CD19⁻ cells. Statistical significance was assessed by Fisher’s one-sided exact test pooled from 3 independent experiments.

**(G)** Histogram showing FasL expression on CAR-T after 48 h co-culture with populations labeled on the right.

**(H)** Histogram showing FasL expression on CD4^+^ and CD8^+^ CAR-T after co-culture with mixed-population CD19^+/^⁻ target cells.

Bars show mean ± s.e.m. Statistical significance was determined by two-way ANOVA with Tukey’s multiple-comparisons test, unless otherwise indicated. *p < 0.03, **p < 0.02, ***p < 0.0002, ****p < 0.0001.

**Supplemental Figure S4. IAPi enhance CAR-T bystander killing without altering tumor growth *in vivo*. Related to Figure 4.**

**(A)** Representative changes in viable cell populations with and without αFas agonist antibody in the presence of various inhibitors of Fas signaling pathway regulators. Percent live in gate is shown in the lower right corner of each plot.

**(B)** Heatmap of extended data in (A).

**(C)** Survival curves for mice in No CAR-T cohorts with and without birinapant treatment, as in Figure 4F. 95% confidence interval represented by shaded area above and below survival lines.

**(D)** Normalized viable counts of CD19⁻Fas^+^ and CD19⁻Fas⁻ tumor cells following 72 h mixed-population co-culture with CAR-T and increasing doses of CUDC-427 (IAPi).

**(E)** Representative changes in populations of mixed-population co-cultures from **(D)**. The numbers in each quadrant are cell counts (x10^3^); bystander killing is highlighted by red boxes.

**(F)** Normalized viable counts of CD19⁻Fas^+^ and CD19⁻Fas⁻ tumor cells following 72 h mixed-population co-culture with CAR-T and increasing doses of xevinapant (IAPi).

**(G)** Representative changes in populations of mixed-population co-cultures from **(F)**. The numbers in each quadrant are cell counts (x10^3^); bystander killing is highlighted by red boxes.

**(H)** Normalized viable counts of CD19⁻Fas^+^ and CD19⁻Fas⁻ OCI-Ly8 tumor cells following single-population co-culture with CAR-T and increasing doses of birinapant.

**(I)** Normalized viable counts of CD19⁻Fas^+^ and CD19⁻Fas⁻ OCI-Ly8 tumor cells following mixed-population co-culture with CAR-T and increasing doses of birinapant.

**(J)** Soluble FasL (sFasL) levels measured by ELISA from the supernatant of mixed-population co-cultures with or without CAR-T in the presence or absence of ADAM10 inhibitor (GI254023X, GIX), as in Figures 4H-J.

Dots indicate replicates; bars show mean ± s.e.m. Statistical significance was determined by two-way ANOVA with Tukey’s multiple comparisons (D, F), Dunnett’s multiple comparisons (J), or by pre-specified two-tailed t-tests (H-I).*p < 0.03, **p < 0.02, ***p < 0.0002, ****p < 0.0001.

**Supplemental Figure S5. CD4^+^ CAR-T can mediate bystander killing and Fas knockout (FasKO) rescues their counts in the presence of IAPi. Related to Figure 6.**

**(A)** Normalized viable counts of CD19^+^Fas^+^, CD19^+^Fas⁻, CD19⁻Fas^+^, and CD19⁻Fas⁻ tumor cells following 48 h mixed-population co-culture with CD4^+^, CD8^+^, or CD3^+^ CAR-T.

**(B)** Normalized viable counts of CD4^+^ and CD8^+^ CAR-T following 48 h mixed-population co-culture in the presence or absence of birinapant (5 µM).

Dots indicate replicates; bars show mean ± s.e.m. Statistical significance was determined by pre-specified t-tests. *p < 0.03, **p < 0.02, ***p < 0.0002, ****p < 0.0001.

**Supplemental Figure S6. Targeting non-tumor cell features to bystander kill cancer cells. Related to Figure 7.**

**(A)** Flow cytometry histogram showing Fas expression on ID8 ovarian cancer cells at baseline and following 24 h stimulation with IFNγ and TNFα.

**(B)** Normalized viable cell counts of FOLR2⁺ Raw264.7 macrophages and FOLR2⁻ B16 melanoma cells after 48 h co-culture with anti-FOLR2 CAR-T in the presence or absence of birinapant (5 µM). Bystander killing of FOLR2⁻ B16 cells is highlighted in red. *n* = 3 technical replicates.

**(C)** Schematic of mouse bispecific antibody (mBsAb)-directed bystander killing. mBsAb binds to primary antibody (1° Ab) targeting CD11b and engages CD3^+^ T cells, leading to Fas-mediated bystander killing of CD11b⁻ cancer cells.

**(D)** Normalized viable cell counts of CD11b⁻ A20 cells after 48 h co-culture with CD3^+^ T cells and αCD11b (1° Ab), mBsAb, or both with or without birinapant (5 µM). *n* = 3 technical replicates.

Dots indicate replicates; bars show mean ± s.e.m. Statistical significance was determined by two-way ANOVA with Tukey’s multiple comparisons. *p < 0.03, **p < 0.02, ***p < 0.0002, ****p < 0.0001.

**Supplemental Table S1. sgRNA sequences used to generate knockout cell lines.** For mouse CIITA, four gRNA were pooled and no single optimal sequence was determined. For Human MS4A1, the two sequences were used interchangeably.

**Supplemental Video S1. Microwell imaging of Fas^+^ bystander killing events**

**Supplemental Video S2. Microwell imaging of Fas**^⁻^ **bystander killing events**

## Resource availability

### Lead Contact

Requests for further information and resources should be directed to and will be fulfilled by the lead contact, Joshua Brody, MD.

### Materials Availability

The cell lines and vectors generated in this study are available from the lead contact with a completed materials transfer agreement.

### Data and Code Availability

Bulk RNA-seq datasets were accessed via dbGaP (phs003145.v1.p1), Kite Pharma, and dbGaP (phs001444.v2.p1).

#### Acknowledgements

The authors would like to thank the Flow Cytometry CoRE at the Icahn School of Medicine at Mount Sinai, BioMedical Engineering and Imaging Institute at the Icahn School of Medicine at Mount Sinai, Microscopy CoRE at the Icahn School of Medicine at Mount Sinai, and the Center for Comparative Medicine and Surgery at the Icahn School of Medicine at Mount Sinai. We are also grateful to the University of Chicago Human Disease and Immunology Core, and the Animal Resource Center for their assistance. We extend our appreciation to Lishi Xie, Ph.D., Xiufen Chen, Ph.D., Sidney Wang, and Alexandra Rojek, M.D. for their invaluable critical feedback throughout the project, and to Elisa Sanchez for assistance with microwell imaging parameter optimization.This research was supported by **. MJ. Lin was supported by NIH T32AI078892 and NIH 5T32GM007280. JK. Chorazeczewski was supported by NIH 5T32AI007090-44, 5T32AI007090-45, and the David and Etta Jonas Center for Cellular Therapy. J Kline received support from the Jonas Center for Cell Therapy Research.

## Author contributions

M.J.L. and J.K.C. conceptualized and performed experiments, validated, analyzed and interpreted data, wrote the original draft, reviewed and edited manuscript. G.P., M.S., S.R., N.H.H., X.X., I.O., R.D. designed and performed experiments. A.C. analyzed clinical data. J.M.T. provided anti-FOLR2 CAR-T viral vector and provided expertise on vector design. D.C. and M.H. assisted with microwell microscopy experimental design. R.U. helped with conceptualization. B.D.B., M.M. helped with conceptualization, supervision, funding acquisition, methodology, reviewing and editing the manuscript. J.K. and J.D.B were responsible for study conceptualization, supervision, project administration, funding acquisition, methodology, writing the original draft, reviewing and editing the manuscript. All authors reviewed and approved the manuscript.

## STAR ⍰ Methods

### Experimental model and study participant details

#### Genetically engineered cell line generation

A20-GFP and -mCherry cells were previously generated by lentiviral transduction.^34^ 20-mCherry cells were spin-transduced with firefly luciferase lentiviral supernatant (LV.PL12.Fluc.T7.PGK.PuroR.SV40A) at MOI 1.6 in 8 µg/mL polybrene and selected with puromycin (2 µg/mL) for 12 days. Cells were subcutaneously injected in Balb/c mice and allowed to grow for 21 days before tumors with the brightest signal were harvested and pooled; this *in vivo* passaging was done three times. Harvested, pooled cells were then clonally selected by single-cell FACS and screened for luciferase expression; 23 positive clonal populations were pooled to reduce clonal heterogeneity.

Knockout (KO) lines were generated by electroporation of Cas9:sgRNA ribonucleoprotein (RNP) complexes. For mouse lines, Cas9-2NLS protein was complexed with sgRNA (molar ratio 1:1.15) for 10 min at room temperature, diluted in 1× TE, mixed with 4 × 10^5^ cells in nucleofector solution (SF Cell Line Solution:Supplement 1, 5:1; Lonza V4XC-2032), and transferred to a 20 µL cuvette. Cells were nucleofected with a Lonza 4D-Nucleofector X (program EH-115 or CM-138) and recovered in 96-well plates for 2 to 3 days. For human lines, RNPs were assembled (sgRNA:Cas9 1.25:1) for 10 min at room temperature; 5 × 10^5^ cells in Neon Resuspension Buffer T were electroporated using the Neon Transfection System (1,400 V, 10 ms, 3 pulses) with 10 µL tips. KO efficiency was assessed by flow cytometry; KO populations were FACS-sorted and re-validated 3 days post-sort prior to expansion and cryopreservation.

Single-gene KOs used sgRNAs listed in STAR Methods Table and Supplementary Table S1. Multi-gene KOs were generated sequentially by nucleofecting a subsequent sgRNA into the parental KO.

#### Animals

Balb/c WT mice (Taconic Biosciences), Balb/c CD45.1 (CByJ.SJL(B6)-Ptprc^a^/J; Jackson Labs, cat no. 006584); and FVB-Tg(CAG-luc,-GFP)L2G85Chco/J (Jackson Labs, cat no. 008450) were housed at the animal facility of the Icahn School of Medicine at Mount Sinai. Balb/c Rag^-/-^ mice (Jackson Labs, strain #003145) and NOD.Cg-*Prkdc^scid^ Il2rg^tm1Wjl^*/SzJ (NSG, Jackson Labs, strain #005557) mice were purchased from the Jackson Laboratory. Balb/c Rag^-/-^ and FVB mice were bred inhouse at the Icahn School of Medicine at Mount Sinai while NSG mice were housed in a specific pathogen-free facility at the University of Chicago. All experiments were conducted in accordance with protocols approved by either the Icahn School of Medicine at Mount Sinai or the University of Chicago’s Institutional Animal Care and Use Committee following guidelines established by the National Institutes of Health.

#### Xenograft mixed-antigen tumor model

Immunodeficient NSG mice were intravenously injected with mixed-antigen OCI-Ly8 lymphoma tumors composed of 80:20% CD19^+^:CD19⁻Fas^+^ or CD19⁻Fas⁻ cells and allowed to establish for 7 days before adoptive transfer of human CAR-T (1.0 x 10^6^ per mouse) by tail-vein injection (Figure 3A). Mice were assessed for survival three times per week and closely monitored for signs of systemic disease (e.g., hind limb paralysis), for which they were euthanized.

#### Syngeneic mixed-antigen tumor models^82^

Mixed-antigen tumors (2.5 x 10^6^ cells per mouse) containing 80:20% CD19^+^:CD19⁻ Fas^+^ or Fas⁻ cells were injected subcutaneously (s.c.) into the flanks of Balb/c mice (Figure 2A). For real-time tracking, 80:20% CD19^+^:CD19⁻Luc^+^ were injected into the right and left flanks of Balbc *Rag1^-/-^* mice (Figure 3A). Bioluminescent imaging (IVIS) was conducted twice per week following injections and caliper measurements were taken as soon as tumors became palpable. Tumor volume (mm^3^) = *L x W x H*, where *L* (mm) is the major axis of the tumor, *W* (mm) is the minor axis perpendicular to *L* in the same plane, and *H* (mm) is the minor axis perpendicular to *L* in the orthogonal plane. When tumors reached ∼500 mm^3^, mice were randomized and conditioned with 4 Gy total body irradiation one day prior to adoptive transfer of CAR-T (7.5 x 10^5^ cells per mouse for non-tracking experiments, 6.0 x 10^6^ cells per mouse for IVIS-tracking experiments) by tail-vein injection.

#### Luciferase^+^ CAR-T tracking in vivo

FVB-Tg(CAG-luc,-GFP)L2G85Chco/J mice were backcrossed with Balb/c CD45.1 mice for 4 generations to create F4. T cells were isolated from the spleens and lymph nodes of F4 and Luc^+^ anti-FOLR2 CAR-T were prepared as described below (See Mouse CAR-T production). Balb/c Rag-/- mice were s.c. injected with 2.5 x 10^6^ A20 tumor cells. When tumors reached ∼250 mm^3^, mice were subjected to 2 Gy total body irradiation one day before intravenous (i.v.) adoptive transfer of 1.5 x 10^6^ Luc^+^ anti-FOLR2 CAR-T. Mice were imaged by IVIS for 5 days post-CAR-T. See Figures 7E-H.

#### Bioluminescent Imaging (IVIS)

Mice were induced in a chamber with 2.5% isoflurane before retro-orbital injection of 7.5 mg/kg D-Luciferin (Revvity, cat no. 122799). Injected mice were transferred to the imaging chamber under 2.5% anesthesia and imaged after 5 min of incubation for 2-60 seconds exposure. Images were collected by the Living Image software (Revvity). Mice were imaged twice per week on Monday and Thursday.

#### Birinapant studies

The same mixed-antigen tumor model described above was used to establish tumors in Balb/c mice. In addition, 2 Gy total body irradiation was given one day prior to adoptive transfer of CAR-T (7.5 x 10^5^ cells per mouse) by tail-vein injection. Birinapant (15 mg/kg) or no drug was injected intraperitoneally into mice starting one day before CAR-T transfer, and dosed twice per week for a total of 10 doses.

### Method details

#### Bulk RNA-seq analysis

Bulk RNA-seq data for the Stanford cohort was obtained via dbGaP (phs003145.v1.p1)^83^ and for ZUMA-1 via Kite Pharma as described in Upadhyay et al.^34^ Bulk RNA-seq data for the NCI cohort was obtained via dbGaP (phs001444.v2.p1), and a count matrix was compiled using STAR count files available from the Genomic Data Commons (GDC). Counts were normalized using the variance-stabilizing transformation (*vst*) in DESeq2 (v1.38.3).^84^ Outliers were assessed by a PCA plot and hierarchical clustering. Duplicate samples for the same subject were summed prior to normalization. To combine the Stanford and ZUMA-1 data sets, batch correction was performed using the *removeBatchEffect* function in limma (v3.54.2).^85^

#### Gene Set Variation Analysis (GSVA)

Gene set scores were quantified using Gene Set Variation Analysis (GSVA) (v1.46.0).^86^ The Death Receptor Signature (DRS) was quantified using genes from Singh et al.^32^

#### Survival analysis of CAR-T patient cohorts

Kaplan-Meier survival analysis was performed using the survival package (v3.4.0) and the survminer package (v0.4.9). The log-rank test was used where indicated. To determine the optimal cut point for survival analysis, the max-stat method was used using the *surv_cutpoint* function.^87^

#### Cell line maintenance

A20, EZB-P6, Raji lymphoma cells, RAW264.7 macrophages, ID8 ovarian cancer cells, and B16 melanoma cells were maintained at 37 °C with 5% CO₂ in RPMI-1640 supplemented with 10% (v/v) FBS, antibiotic-antimycotic (100 U/mL penicillin, 100 µg/mL streptomycin, 0.25 µg/mL amphotericin B), and 50 µM 2-mercaptoethanol. OCI-Ly8 cells were cultured under the same conditions with additional 1% (v/v) MEM Non-Essential Amino Acids and 1% (v/v) HEPES. Suspension lines were passaged every 2– 3 days to maintain 0.2–1.0 × 10^6^ cells/mL; adherent lines were maintained at 60-70% confluence. EZB-P6 cells were a gift from the Béguelin lab (Cornell). RAW264.7 and ID8 cells were gifts from the Brown lab (Mount Sinai). All lines were authenticated by short tandem repeat (STR) profiling and tested mycoplasma-negative by PCR prior to use and at regular intervals. Master stocks were cryopreserved in FBS + 10% DMSO and stored in liquid nitrogen; working banks were thawed and used for ≤3 weeks post-thaw.

#### Flow cytometry and cell sorting

Cells were resuspended in FACS staining buffer (Hank’s Balanced Saline Solution (HBSS) + 0.5% BSA, 0.1% sodium-azide, 2 mM EDTA). The following mouse antibodies were used: CD3-APC, CD4-BV785, CD45.1-BV421, CD45.1-AF700, CD8a-BV711, Fas-PE-Cy7, Fas-APC, Fas-PerCP-Cy5.5, B220-PerCP-Cy5.5, TCRβ-BV650, FasL-PE, cleaved caspase-3**. The following human antibodies were used: CD3, TCRβ, CD4, CD8, CD19, CD20, Fas, FasL**. CAR expression was detected using GFP. For quantification of absolute cell numbers (tumor or T cells) acquired during flow cytometry, a homogeneous suspension of Precision Count Beads (Biolegend, cat no. 424902) uniformly added to each sample prior to acquisition; counts were normalized to the beads acquired per sample. Cell viability was assessed with Fixable Viability Dye eFluor 780 (ThermoFisher, cat no. 65-0865-14) or Annexin V (Biolegend, 640906) according to manufacturer’s protocol. Unless otherwise specified, all gating strategies followed: FSC-A vs. SSC-A (cells), FSC-W vs. FSC-H (singlets 1), SSC-W vs. FSC-A (singlets 2), FSC-A vs. Live/Dead (Viability). Data were acquired on Attune NxT (ThermoFisher) or BD LSRFortessa (BD Biosciences) and analyzed with Cytobank (Cytobank, Inc., Santa Clara, CA; https://www.cytobank.org). Cell sorting was completed with FACSAria (BD Biosciences) or CytoFLEX SRT (Beckman Coulter).

#### CAR vector cloning

All CAR constructs were expressed an MSGV backbone and composed of an scFv, CD28 costimulatory domain, and CD3ζ domain with 1^st^ and 3^rd^ ITAM domain deletions (1-3mut). For anti-CD19 CAR, the MSGV1-1D3-28z.1-3mut vector was used (Addgene #107227). Cloning of the WTFasL.CAR and mutFasL.CAR was completed by stepwise restriction enzyme digest and ligation (EcoRI-HF, NEB #R3101S; SalI-HF, NEB #R3138S) from sequences by Twist Bioscience, CA, USA. WTFasL and mutFasL were both preceded by a P2A self-cleaving sequence and followed by T2A and GFP. Fully cloned CAR constructs were verified by sequencing. For anti-FOLR2 CAR, the CL10 scFv^88^ was cloned into an MSGV-CD28-CD3ζ all ITAM intact backbone (Addgene #107226).

#### Retrovirus production

Phoenix-Ecotropic (PE) cells were maintained in complete Iscove’s Modified Dulbecco’s Medium (IMDM) in 15 cm petri dishes at 37°C with 5% CO2. One day prior to transfection, cells were plated at 9.0 x 10^6^ per dish. The next day, fresh IMDM media was exchanged 3 hours prior to transfection. Cells were transfected via the calcium-phosphate DNA precipitation method. Briefly, 2 mM CaCl2, 0.1X TE buffer, and plasmid of interest were mixed at RT and incubated for 5 min. 2X HBS was added dropwise to DNA mixture while vortexing to form DNA-CaPi precipitate and incubated at RT for 10 min to allow for crystal formation. DNA solution was added on top of PE cells. 14-16 h later, media was exchanged for fresh IMDM. 24 h later, RPMI was exchanged. 24 h later, viral supernatant was collected, filtered through a 0.2 mm disc filter, aliquoted, and stored at -80°C for later use.

#### Lentivirus production

Lenti-X™ 293T cells (293T cells) were maintained in 60mm cell culture dishes with 10% FBS in DMEM at 37°C with 5% CO_2_. 2.0-3.0 x 10^6^ cells were plated per dish 1 day prior to transfection. Fresh media was added to the dish 1 to 3 h prior to transfection. Cells were transfected using a polyethylenimine (PEI)-mediated transfection protocol. Briefly, the plasmid of interest, packaging plasmid (pCMV-dR8.2-dvpr), envelope plasmid (pMD2.G), and PEI are vortexed and incubated RT for 10 minutes. the DNA-PEI mixture is then added dropwise to the 293T cells. After 48 hours, media was harvested from the 293T cells and spun for 1000 x g for 10 minutes or ultracentrifuged (Sorvall WX 100+, SW-28 rotor) at 20,000 rpm (∼40,000 x g) for 2.5 h at room temperature to pellet any remaining cells while the supernatant was collected, filtered through a 0.2 mm disc filter, aliquoted and stored at -80°C.

#### Mouse CAR-T production

CD3^+^ cells from spleens and lymph nodes of 8 to 12-week-old Balb/c CD45.1 mice were negatively selected by Invitrogen Untouched Mouse T cell kits (ThermoFisher, cat no. 11413D) and activated with Mouse T-activator CD3/28 Dynabeads (ThermoFisher, cat no. 11453D) in 100 IU/mL IL-2 (PeproTech, #212-12-1MG) in RPMI media. One day after activation, cells were spin-transduced in non-tissue culture-treated 6-well plates coated overnight at 4°C with 20 ng/mL RetroNectin® (Takara, cat no. T101) with 1 mL of retrovirus supernatant for 1 h at 2000 x g in 30°C. After 2 days of CD3/28 activation, Dynabeads were removed by magnet column. Transduced cells were replated at 0.8-1.2 x 10^6^ cells/mL in 50 IU/mL IL-2 every 24 h. Expanded cells were used 3-7 days after transduction, with no additional IL-2 added into co-culture wells. For repeat *in vitro* testing, large batches of CAR-T were cryopreserved in CryoSFM media (Millipore Sigma, cat no. C-29912) and kept at -80°C. Cells were thawed 2 days prior to plating and rested in RPMI supplemented with 100 IU/mL IL-2. The viability of cells was increased by Ficoll-gradient centrifugation 1 day after thawing.

#### Human CAR-T production

Human peripheral blood mononuclear cells (PBMCs) were isolated from whole blood of healthy donors by density gradient centrifugation using Ficoll-Paque PLUS. Isolated PBMCs were rested overnight in T cell expansion media (TexMACS™ GMP Medium) + 10% FBS + IL-7 and IL-15 (5 ng/mL) after which adherent monocytes were removed and remaining non-adherent lymphocyte population were activated with ImmunoCult™ Human CD3/CD28/CD2 T Cell Activator. One day after activation, 5.0 x 10^4^ CD3^+^ T cells were seeded in 100 µL of T cell expansion media in 96-well non-tissue culture-treated round bottom plates. T cells were spin-transduced with lentivirus supernatant (MOI 10) and protamine sulfate (10 µg/mL) at 900 x g for 1 h at 32°C. Transduced cells were kept at 0.5-2.0 x 10^6^ cells/mL during expansion in IL-7 and IL-15 in T cell expansion media and used 9 to 11 days post-transduction. Large batch CAR-T cells were cryopreserved in freezing media (10% DMSO in FBS) and kept at -80°C. Cells were rested for 1 to 2 days prior to use.

#### T cell isolation for BsAb assays

Mouse CD3^+^ T cells were isolated from the spleens and lymph nodes of mice as described above. CD3/28 Dynabead activation beads were removed by magnet column after 2 days, and cells were expanded in RPMI + 50 IU/mL IL-2 for 3-5 days until co-culture with target cells and redirecting antibodies.

Human CD3^+^ T cells were isolated with the MojoSort Human CD3 T Cell Isolation Kit (Biolegend, cat no. 480022) from PBMC collected by density gradient centrifugation. Isolated cells were plated with target populations at indicated scales and with indicated doses of CD3xCD20 BsAb for 48-72 h.

#### IAP inhibitors

The IAP inhibitor birinapant (TL32711, MedChemExpress, cat no. HY-16591) was resuspended in DMSO for a stock solution of 10mM. The IAP inhibitor CUDC-427 (GDC-0917, MedChemExpress, cat no. HY-15835) was received reconstituted in DMSO at a stock solution of 10mM. The IAP inhibitor xevinapant (AT406, Selleckchem, cat no. S2754) was resuspended in DMSO for a stock solution of 10mM. For *in vivo* administration, birinapant was dissolved in the vehicle solution 12.5% (w/v) Captisol (MedChemExpress, cat no. HY-172405) in ddH_2_O and slowly titrated to a pH 4.0 with HCl and NaHCO_3_, and i.p. injected at 15 mg/kg twice per week for a total of 10 doses.

#### In vitro killing assays

##### CAR-T bystander killing assays^82^

Either murine A20 or human OCI-Ly8 target cells were mixed as single antigen co-cultures (CD19^+^Fas^+/^, or CD19⁻Fas^+/^) or mixed antigen co-cultures (CD19^+^Fas^+^, CD19^+^ Fas⁻, CD19⁻Fas^+^, CD19⁻ Fas⁻) in a 1:1 ratio of 2.5-5.0 x 10^4^ cells each per well in U-bottom 96-well plates (Fisher Scientific, cat no. 08-772-17). CAR-T were added to each co-culture at an effector-to-target (E:T) ratio of 2:1 (1.0-2.0 x 10^5^ CAR-T) or 8:1 for FasKO experiments. For drug treatment conditions, a 3X solution was prepared and 80 µL added to each well such that the final volume was 240 µL. After indicated co-culture times (48-72 h), cells were collected for flow cytometry, or supernatant for ELISA. Mouse samples were stained in FACS buffer for 20 min at either 37°C or 25°C, while human samples were stained for 20 min at 4°C before fixation in IC Fixation buffer (ThermoFisher). For intracellular stains, samples were permeabilized and stained for 30 min at 25°C.

Cell counts for each population were normalized to the No CAR-T control condition. Bystander killing index was calculated as 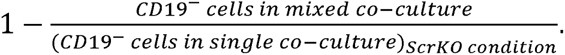

##### Transwell CAR-T bystander killing assays

Transwell killing assays were conducted as described above, with the indicated populations (1.0 x 10^4^ cells per population in each compartment) seeded inside (within) and/or outside (below) a transwell insert separated by a permeable membrane with 0.4 µm pores (Fisher Scientific, cat no. 08-770) within a 24-well plate. After 48 h, the cells from inside (within) and outside (below) the transwell were separately transferred to a 96-well plate for staining and acquisition for flow cytometry. See Figure 2H.

##### Blocking antibody killing assays

Co-cultures were conducted as described above, with the addition of blocking antibodies at a final concentration of 100 µg/mL: Isotype (Rat IgG1, kappa, BioXCell, cat no. BE0088), anti-IFNγ (R4-6A2, BioXCell, cat no. BE0054), TNFα (XT3.11, BioXCell, cat no. BE0058), anti-TRAIL-R (MD5-1, BioXCell, cat no. BE0161), FasL (MFL3, BioXCell, cat no. BE0319). Samples were washed 3 times with HBSS prior to staining according to protocol described above. See Supplemental Figure S4G.

##### Fas agonist killing assays

Mouse or human lymphoma cell lines were seeded in 96-well plates at 5.0 x 10^4^ cells/well and incubated for 48-72 h with anti-Fas agonist antibody at indicated concentrations (mouse: Jo2, BD Bioscience, cat no. 554255; human: CH11, Fisher Scientific, cat no. 05-201-MI). Cell counts and viability were assessed by flow cytometry as described above.

##### Human BsAb-directed bystander killing assays

Human OCI-Ly8 or Raji target cells were co-cultured as mixed-antigen populations (CD20^+^, CD20⁻Fas^+^, CD20⁻Fas⁻) in a 1:1 ratio of 2.5 x 10^5^ cells each per well in a U-bottom 96-well plate with 2:1 E:T pre-activated CD3^+^ T cells in the absence or presence of CD20xCD3 BsAb at indicated doses (Epcoritamab, GEN3013, MedChemExpress, cat no. HY-P99931). For additional birinapant co-culture, a 3X drug solution was prepared and 80 µL added to indicated wells such that the final volume was 240 µL in each well. Samples were analyzed after 48-72 h co-culture by flow cytometry as described above.

##### RAW264.7 macrophage Bystander killing Assays

Adherent cells (RAW264.7, ID8, or B16) were seeded in 96-well flat-bottom plates one day prior to addition of CAR-T at a density of 1.0 x 10^4^ cells per well per population. A20 bystander cells were seeded at 2.0 x 10^4^ the same day as CAR-T. When Fas^+/^ bystander cell populations were used, 0.5 x 10^4^ of each respective adherent population and 1.0 x 10^4^ of each respective suspension population was added to the mixture such that the total number of bystander tumor cells remained at the same number as the on-target RAW264.7 population (1:1:2, effectively 1:1). 2:1 E:T ratio of anti-FOLR2 CAR-T was plated one day following adherent cell line seeding and co-cultured for 48 h. For additional birinapant co-culture, a 3X drug solution was prepared and 80 µL added to indicated wells such that the final volume was 240 µL in each well. For staining and sample preparation for flow cytometry, non-adherent cells were first transferred to a separate 96-well plate. Adherent cells were then washed 2 times with HBSS before detaching with EDTA solution (FACS buffer + 0.5% EDTA) and combining with the suspension cells in the 96-well plate from each respective well. Cells were stained and analyzed as described above.

##### mBsAb-directed Bystander Killing Assays

RAW264.7 cells were seeded in 96-well flat-bottom plates one day prior to addition of CAR-T at a density of 1.0 x 10^4^ cells per well. The following day, A20 cells were seeded on top at 1.0 x 10^4^ cells per well. 2:1 E:T pre-activated CD3^+^ T cells were added in culture with and without the following reagents: anti-CD11b antibody (primary 1° Ab, 10nM), mBsAb (10nM), or both. For additional birinapant co-culture, a 3X drug solution was prepared and 80 µL added to indicated wells such that the final volume was 240 µL in each well. See Supplemental Figure S6D.

#### Microwell Imaging Assays ^82^

PDMS grids with 50 x 50 x 25 µm wells were adhered to the bottom of an 8-chamber glass slide and submerged in HBSS overnight at room temperature. Chambers were washed with HBSS 2x and 200µL of 10µg/mL fibronectin solution was added to each chamber and incubated at 37°C for 1 h. CD19^+^ cells were labeled with CellTrace CFSE at 1:2000 dilution for 5 min at 37°C, CD19⁻ cells were labeled with CellTrace Violet at 1:2000 dilution for 5 min at 37°C, and CAR-T were labeled with CellVue Maroon at 20 µM dilution for 5 min at 37°C. Cells were quenched with media and washed twice with media before resuspension at 4.0 x 10^4^ cells/mL. Chambers were aspirated after fibronectin coating and seeded with 200 µL of a 1:1 mixture of CFSE-labeled on-target cells and CT-Violet-labeled bystander cells, followed by centrifugation at 1500 rpm for 5 min. Grids were assessed under light microscope to confirm a density of 1-3 cells per microwell. Additional cell suspension was added as needed to increase seeding, with excess media carefully removed after each spin. CAR-T suspension was added on top, centrifuged, and adjusted until each well contained 3-6 cells. The final chamber volume after cell seeding was 200 µL.

Chambers were imaged on a Leica DMi8 widefield microscope equipped with an incubation chamber. Images were acquired every 5-10 min from regions spanning the majority of the grids, in brightfield and fluorescence channels (CFSE: 498/517 nm, CT Violet: 405/450 nm, PI: 535/617 nm, and CV Maroon: 647/667 nm).

Raw Leica files were converted to TIFF file format using FIJI (ImageJ). TIFF files were then analyzed in FIJI by identifying wells-of-interest; wells that were not visible for the full imaging duration or lacking all three cell types were excluded from analysis. Wells-of-interest were visually assessed for cell death events defined as either propidium-iodide (PI) uptake or characteristic morphological changes (membrane blebbing, cell shrinkage). Killing events were categorized as (1) on-target killing, (2) bystander killing, or (3) no killing. Triad-based killing was defined as bystander killing occurring in the context of simultaneous physical contact between CAR-T, on-target, and bystander tumor cells. Representative wells were selected for further image stabilization and visual stitching in DaVinci Resolve 18.6 (Blackmagic Design). No alterations were made to image color, contrast, brightness, or sharpness at this stage.

#### Western blot analysis

Equal numbers of live cells were washed with PBS, lysed using RIPA buffer (Merck, #20-188) with protease-phosphatase inhibitors (Thermo Fisher, #A32961). Equal protein amounts were run on 10–12% SDS-PAGE gels and transferred to 0.2 or 0.45 μm PVDF membranes. Membranes were incubated with anti-FasL antibody (Cell Signaling, #72062S) and detected using Clarity ECL substrate and ChemiDoc MP (Bio-Rad). Protein levels were normalized to β-Actin (Santa Cruz, #sc-47778) and control groups.

#### Enzyme-linked immunosorbent assay (ELISA)

CAR-T were co-cultured with target cells (CD19+, CD19-, or mixed) at 2:1 E:T for 24 and 48 h in standard media. Supernatant was collected and stored -20°C if not used immediately. Analyses were performed using Mouse Fas Ligand/TNFSF6 Quantikine ELISA kits (R&D Biosciences, #MFL000).

#### FasKO CAR-T cells^89^

Mouse CAR-T were generated as described above. On day 3 of expansion, CAR-T were subject to Cas9/sgRNA RNP electroporation using the P3 Primary Cell 4D-Nucleofector X Kit and 4D-Nucleofector Core Unit (Lonza) as described. Scrambled gRNA control (Synthego) or Mouse Fas gRNA (listed in Supplemental Table S1) were used to knockout genes in activated CAR-T. Immediately after electroporation, cells were rested in RPMI without antibiotics for 30 min before being centrifuged and transferred to complete RPMI + 50 IU/mL IL-2. One day after electroporation, dead cells were removed by MACS Dead Cell Removal Kit (Miltenyi Biotec, cat no. 130-090-101). Two days after electroporation, KO efficiency was determined by flow cytometry. Three days after electroporation, CAR-T were collected for co-culture.

### Quantification and statistical analysis

All statistical data analyses, unless detailed in figure legends and Method details, were performed with GraphPad Prism 9.5.1 (GraphPad Software, La Jolla, USA) or higher. Comparisons between three or more groups were performed using two-way ANOVA with Tukey’s multiple comparisons, unless otherwise specified. Data are represented as mean ± s.e.m. unless otherwise noted. Survival data were analyzed using log-rank tests. Sample sizes and replicate numbers are indicated in the main text, figures, or figure legends.

