## Supplemental Figures for "Potentiating CAR-T bystander killing by enhanced Fas/FasL signaling mitigates antigen escape in heterogeneous tumors"

**Supplemental Video S1. Microwell imaging of $Fas^+$ bystander killing events**
**Supplemental Video S2. Microwell imaging of $Fas^-$ bystander killing events**

#### Supplemental Figures

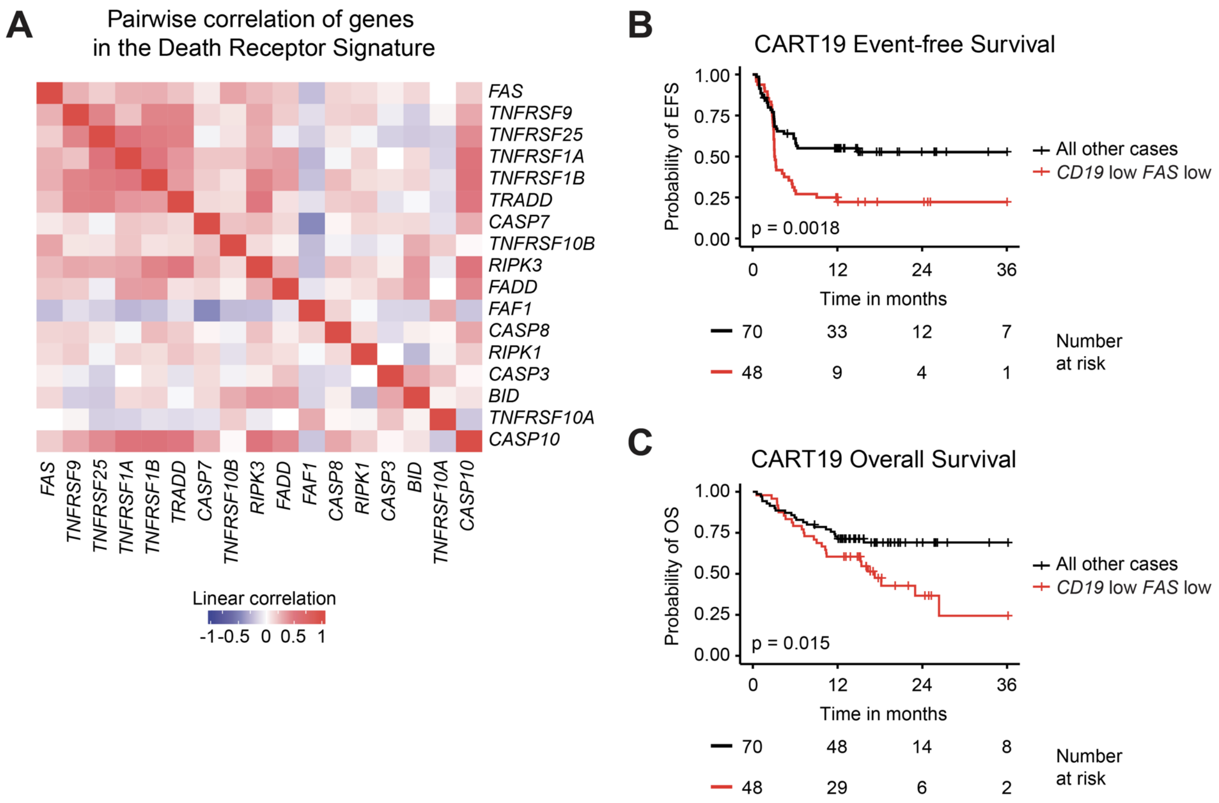

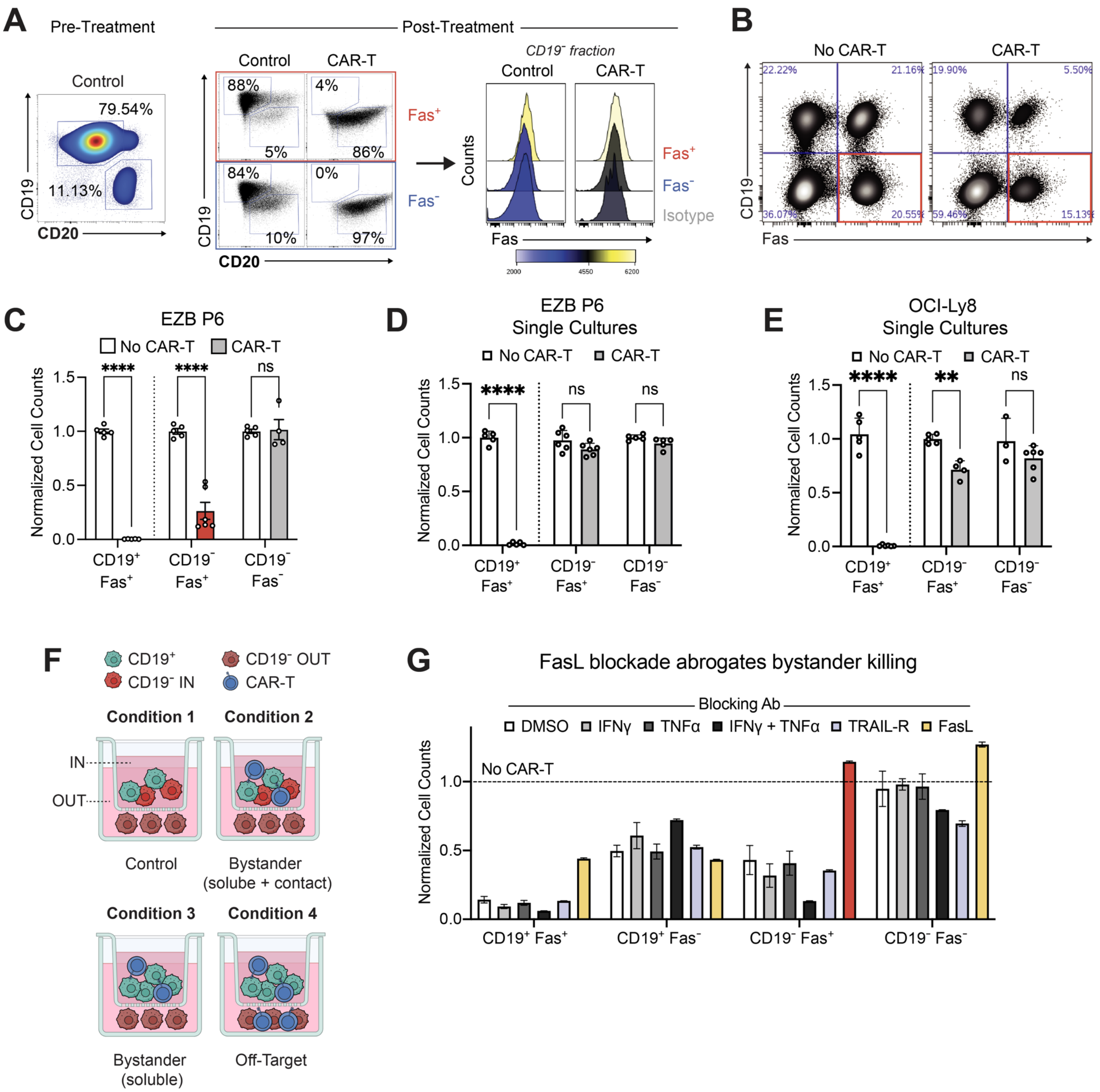
A
Pre-Treatment
Post-Treatment
Control
79.54%
11.13%
CD19
CD20
Control
CAR-T
88%
4%
5%
86%
84%
0%
10%
97%
Fas+
Fas-
CD19⁻ fraction
Control
CAR-T
Counts
Fas
Isotype
B
No CAR-T
CAR-T
22.22%
21.16%
19.90%
5.50%
36.07%
20.55%
59.46%
15.13%
CD19
Fas
C
EZB P6
No CAR-T
CAR-T
Normalized Cell Counts
****
****
ns
CD19+ Fas+
CD19- Fas+
CD19- Fas-
D
EZB P6
Single Cultures
E
OCI-Ly8
Single Cultures
**
F
CD19+
CD19- OUT
CD19- IN
CAR-T
Condition 1
Condition 2
IN
OUT
Control
Bystander
(solube + contact)
Condition 3
Condition 4
Bystander
(soluble)
Off-Target
G
FasL blockade abrogates bystander killing
Blocking Ab
DMSO
IFNγ
TNFα
IFNγ + TNFα
TRAIL-R
FasL
No CAR-T
Normalized Cell Counts
CD19+ Fas+
CD19+ Fas-
CD19- Fas+
CD19- Fas-

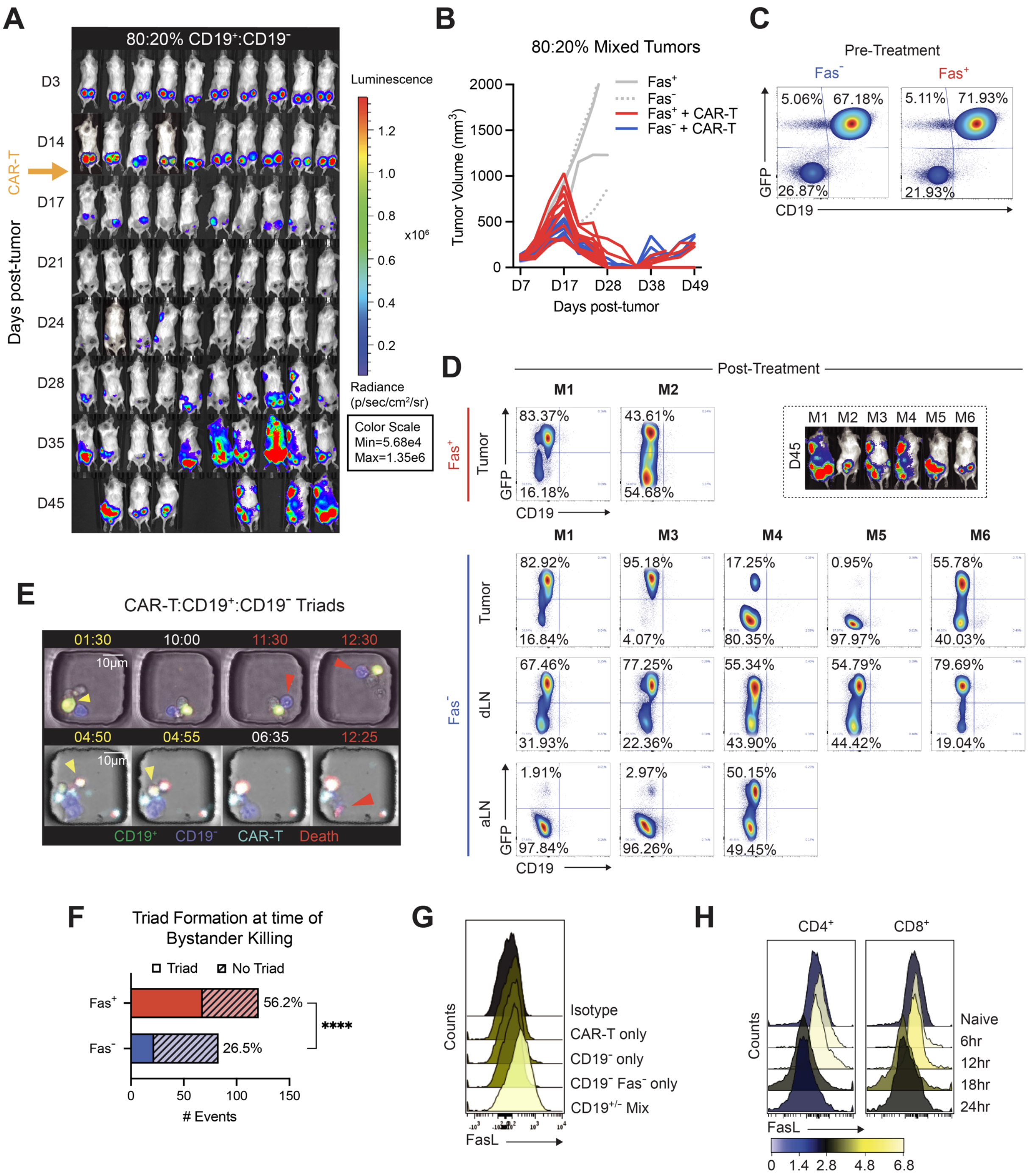
A
80:20% CD19+:CD19-
D3
D14
D17
D21
D24
D28
D35
D45
CAR-T
Days post-tumor
Luminescence
x10^6
Radiance (p/sec/cm2/sr)
Color Scale Min=5.68e4 Max=1.35e6
B
80:20% Mixed Tumors
Fas+
Fas-
Fas+ + CAR-T
Fas- + CAR-T
Tumor Volume (mm3)
Days post-tumor
C
Pre-Treatment
Fas-
Fas+
5.06% 67.18%
26.87%
5.11% 71.93%
21.93%
GFP
CD19
D
Post-Treatment
M1 M2
Fas+
Tumor
83.37% 16.18%
43.61% 54.68%
D45
M1 M2 M3 M4 M5 M6
Fas-
Tumor
82.92% 16.84%
95.18% 4.07%
17.25% 80.35%
0.95% 97.97%
55.78% 40.03%
dLN
67.46% 31.93%
77.25% 22.36%
55.34% 43.90%
54.79% 44.42%
79.69% 19.04%
aLN
1.91% 97.84%
2.97% 96.26%
50.15% 49.45%
E
CAR-T:CD19+:CD19- Triads
01:30 10:00 11:30 12:30
04:50 04:55 06:35 12:25
10μm
CD19+ CD19- CAR-T Death
F
Triad Formation at time of Bystander Killing
Triad
No Triad
Fas+ 56.2%
Fas- 26.5%
****
### Events
G
Counts
Isotype
CAR-T only
CD19- only
CD19- Fas- only
CD19+/- Mix
FasL
H
CD4+
CD8+
Naive
6hr
12hr
18hr
24hr
Counts
FasL

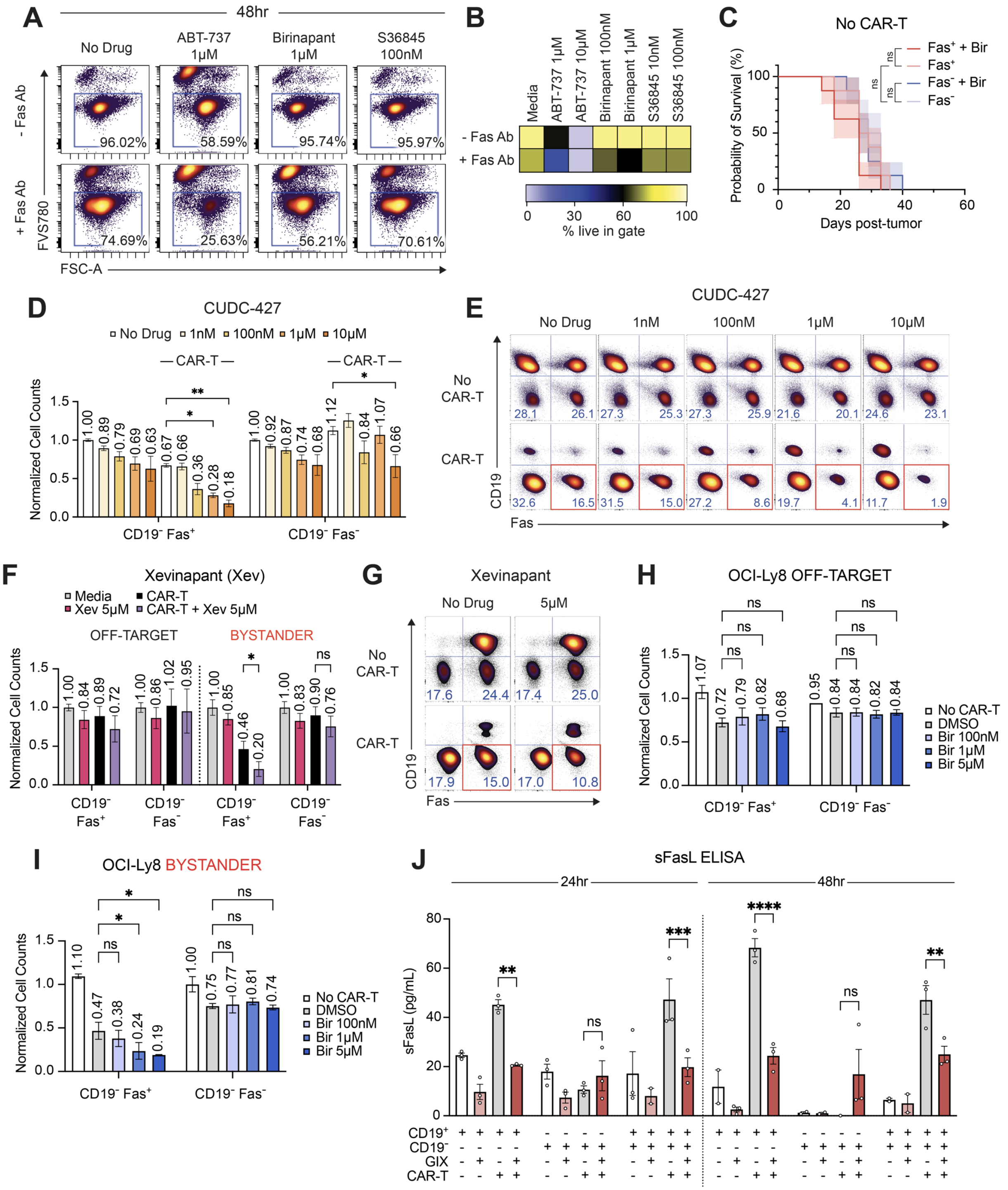

A
48hr
No Drug
ABT-737 1μM
Birinapant 1μM
S36845 100nM
- Fas Ab
+ Fas Ab
FVS780
FSC-A
96.02%
58.59%
95.74%
95.97%
74.69%
25.63%
56.21%
70.61%
B
Media
ABT-737 1μM
ABT-737 10μM
Birinapant 100nM
Birinapant 1μM
S36845 10nM
S36845 100nM
- Fas Ab
+ Fas Ab
% live in gate
C
No CAR-T
Fas+ + Bir
Fas+
Fas- + Bir
Fas-
Probability of Survival (%)
Days post-tumor
D
CUDC-427
No Drug 1nM 100nM 1μM 10μM
CAR-T
Normalized Cell Counts
CD19- Fas+
CD19- Fas-
E
CUDC-427
No Drug 1nM 100nM 1μM 10μM
No CAR-T
CAR-T
CD19
Fas
F
Xevinapant (Xev)
Media
CAR-T
Xev 5μM
CAR-T + Xev 5μM
OFF-TARGET
BYSTANDER
G
Xevinapant
No Drug
5μM
No CAR-T
CAR-T
H
OCI-Ly8 OFF-TARGET
No CAR-T
DMSO
Bir 100nM
Bir 1μM
Bir 5μM
I
OCI-Ly8 BYSTANDER
J
sFasL ELISA
24hr
48hr
sFasL (pg/mL)
CD19+
CD19-
GIX
CAR-T

**A**

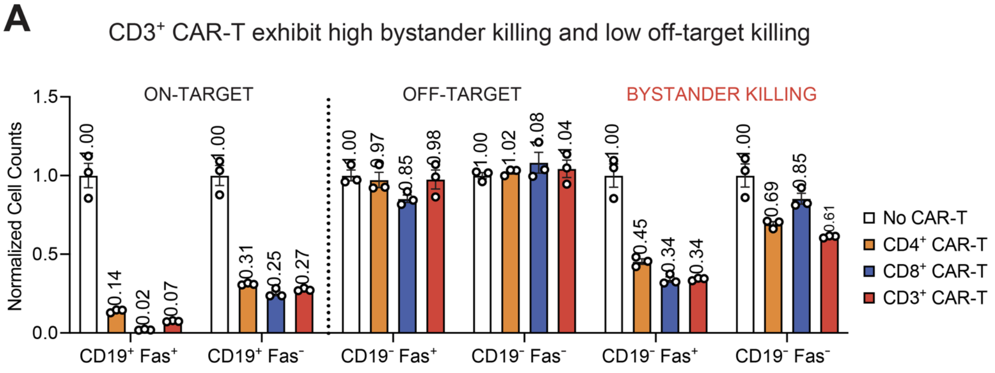
CD3+ CAR-T exhibit high bystander killing and low off-target killing
ON-TARGET
OFF-TARGET
BYSTANDER KILLING
Normalized Cell Counts
1.5
1.0
0.5
0.0
1.00
0.14
0.02
0.07
1.00
0.31
0.25
0.27
1.00
0.97
0.85
0.98
1.00
1.02
1.08
1.04
1.00
0.45
0.34
0.34
1.00
0.69
0.85
0.61
CD19+ Fas+
CD19+ Fas-
CD19- Fas+
CD19- Fas-
CD19- Fas+
CD19- Fas-
No CAR-T
CD4+ CAR-T
CD8+ CAR-T
CD3+ CAR-T

**B**

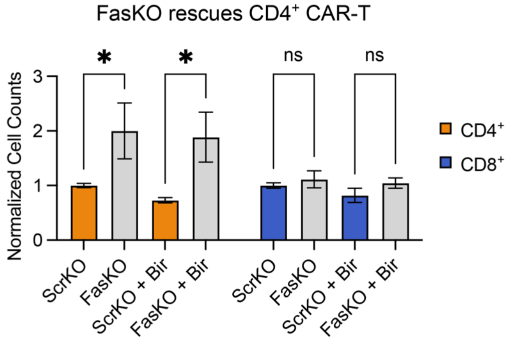
FasKO rescues CD4+ CAR-T
Normalized Cell Counts
3
2
1
0
*
*
ns
ns
ScrKO
FasKO
ScrKO + Bir
FasKO + Bir
ScrKO
FasKO
ScrKO + Bir
FasKO + Bir
CD4+
CD8+

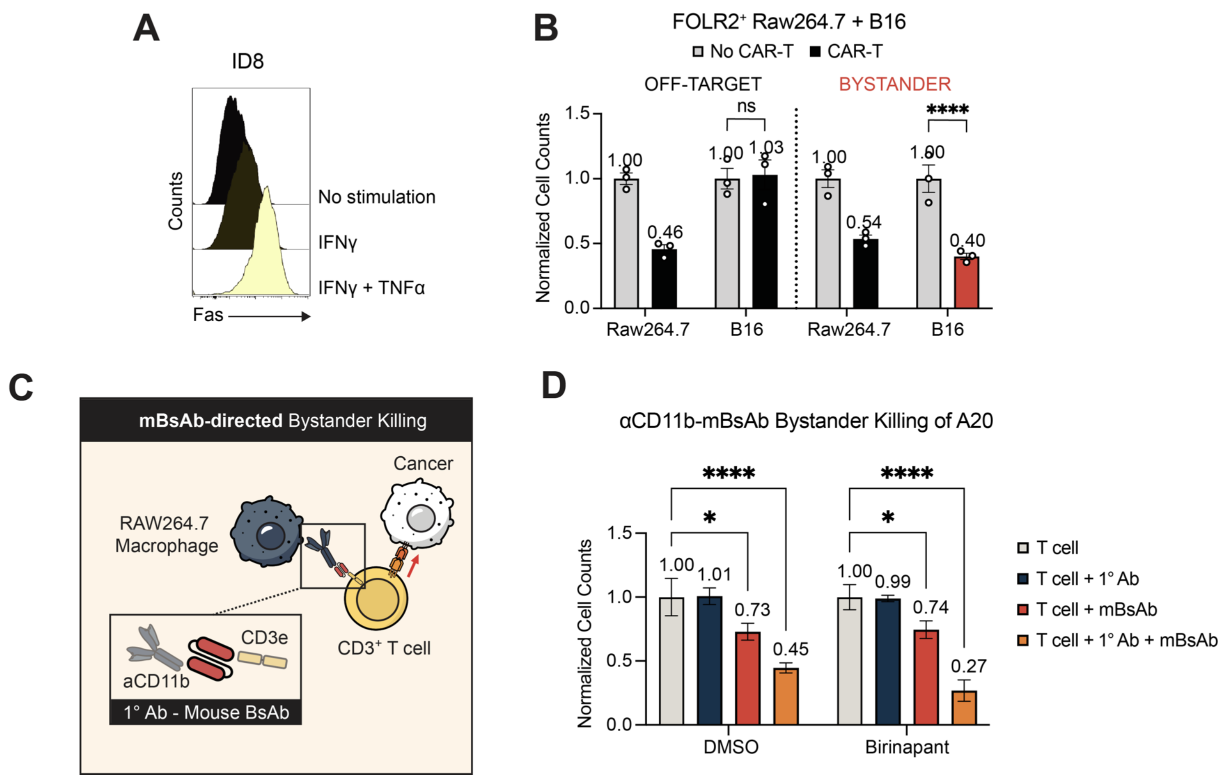


**Supplemental Table S1. sgRNA sequences used to generate knockout cell lines.** For mouse CIITA, four gRNA were pooled and no single optimal sequence was determined. For Human MS4A1, the two sequences were used interchangeably.

| Target | gRNA Sequence (5'-3') | Exon |
|---|---|---|
| Mouse B2M | CACGCCACCCACCGGAGAAU | 2 |
| Mouse CIITA | 1) AACUAUUUGGAUCUCCCCAC,<br>2) CUUCCCGGAAGAGCAUGCUA,<br>3) CCUCUAUGACCAGAUGGACC,<br>4) AGUUCCAGGUAGCUGCCCUC | 7, 6, 2, 2 |
| Mouse CD19 | AGACAGGUGAGGAGUCCGGG | 2 |
| Mouse MS4A1 | GGCCUCUUCCAUAUUACCCU | 3 |
| Mouse FAS | GGCGUCCCAAAGCUUACCAG | 1 |
| Human B2M | GAGUAGCGCGAGCACAGCUA | 1 |
| Human CIITA | GAUAUUGGCAUAAGCCUCCC | 3 |
| Human CD19 | CCUUCUUAAAACUCAGCCUG | 2 |
| Human MS4A1 | 1) CCAUAAUGCCUCCCCAGAGA,<br>2) GGAUCAUCAGAAGACCCCCC | 2, 4 |
| Human FAS | GGAGUUGAUGUCAGUCACUU | 2 |
